# *In silico, In vitro* Screening of Plant Extracts for Anti-SARS-CoV-2 Activity and Evaluation of Their Acute and Sub-Acute Toxicity

**DOI:** 10.1101/2021.09.07.459230

**Authors:** Latha Damle, Hrishikesh Damle, Shiban Ganju, C Chandrashekar, BR Bharath

**Affiliations:** Computational Biology, Atrimed Biotech LLP, Banglore, 560100, India; Atrimed Pharmaceuticals Pvt. Ltd, Banglore, 560001, India

**Keywords:** Molecular docking, Plant extracts, Cytotoxicity, Anti-SARS-CoV-2, *In vivo* toxicity

## Abstract

**Background:** In the absence of a specific drug for COVID 19, treatment with plant extracts could be an option worthy of further investigation.

**Purpose:** To screen the phytochemicals for Anti-SARS-CoV-2 *in silico* and evaluate their safety and efficacy *in vitro* and *in vivo*.

**Method:** The phytochemicals for Anti-SARS-CoV-2 were screened *in silico* using molecular docking. The hits generated from *in silico* screening were subjected for extraction, isolation and purification. The anti-SARS-CoV-2 activity of plant extracts of *Z. piperitum* (ATRI-CoV-E1), *W. somnifera* (ATRI-CoV-E2), *C. inophyllum* (ATRI-CoV-E3), *A. paniculata* (ATRI-CoV-E4), and *C. Asiatica* (ATRI-CoV-E5). The *in vitro* safety and anti-SARS-CoV-2 activity of plant extracts were performed in VeroE6 cells using Remdesivir as positive control. The acute and sub-acute toxicity study was performed in Wistar male and female rats.

**Results:** The percentage of cell viability for ATRI-COV-E4, ATRI-COV-E5 and ATRI-COV-E2 treated VeroE6 cells were remarkably good on the 24th and 48th hour of treatment. The *in vitro* anti-SARS-CoV-2 activity of ATRI-COV-E4, ATRI-COV-E5 and ATRI-COV-E2 were significant for both E gene and N gene. The percentage of SARS-CoV-2 inhibition for ATRI-COV-E4 was better than Remdesivir. For E gene and N gene, Remdesivir showed IC_50_ of 0.15 µM and 0.11 µM respectively, For E gene and N gene, ATRI-CoV-E4 showed IC_50_ of 1.18 µg and 1.16 µg respectively. Taking the clue from *in vitro* findings, the plant extracts *A. paniculata* (ATRI-COV-E4), *W. somnifera* extract (ATRI-COV-E5) and *C. asiatica* extract (ATRI-COV-E2) were combined (ATRICOV 452) and evaluated for acute and sub-acute toxicity in Wistar male and female rats. No statistically significant difference in haematological, biochemical and histopathological parameters were noticed.

**Conclusion:** The study demonstrated the Anti-SARS-CoV-2 activity *in vitro* and safety of plant extracts in both *in vitro* and *in vivo* experimental conditions.

## 1. **Introduction**

Currently, there is no specific drug for SARS-CoV-2 infection. Efforts are being made globally to identify agents for preventive, supportive and therapeutic care. Many plants used in traditional medicine have been reported to have antiviral properties. In the present study, we evaluated cytotoxicity and antiviral activity against SARS-CoV-2 of *Zanthoxylum piperitum* (ZP), *Withania Somnifera (WS), Calophyllum inophyllum* (CI), *Andrographis paniculata, (AP) and Centella Asiatica (CA)*,. We also studied animal toxicity for AP, WS and CA.

ZP leaf extracts have been investigated against influenza viruses, A/WS/33, A/PR/8 and B/Lee/40 and reported to have antiviral activity at low concentration (Choi et al. 2008). ZP extracts have shown inhibition of human rhinovirus 2, human rhinovirus 3, coxsackie A16 virus, coxsackie B3 virus, coxsackie B4 virus, and enterovirus 71 virus (Choi et al. 2016).

WS *has been* extensively studied for its antioxidant, antibacterial, anti-fungal, anti-inflammatory, and hematopoietic activity. A prospective, randomized double-blind, placebo-controlled study of WS root extract demonstrated its safety and efficacy in reducing stress and anxiety in adults (Chandrasekhar et al. 2012). WS contains pharmacologically active compounds like sitoindosides and alkaloids which are reported for various biological activities (Bhattacharya and Muruganandam 2003; Kulkarni & Dhir 2008; Singh et al. 2010; Provino 2010; Sharma et al. 2011). WS has been screened for antiviral activity against Infectious Bursal Disease Virus Replication. The hydro-alcoholic extract of WS roots showed 99.9% inhibition of the virus at its highest nontoxic concentration (Pant et al. 2012).

CA has been used in the Ayurvedic tradition of India and listed in an ancient Indian medical text called ‘Sushruta Samhita’ (Chopra et al. 1986; Diwan et al. 1991). CA is also used in Indonesian islands. The primary active constituents of CA are triterpenoids, which include asiaticosides, madecassoside and madasiatic acid (Singh & Rastogi 1961). CA in combination with Madura cochinchinensis (Lour.) Corner, and Mangifera indica was evaluated for anti-herpes simplex virus (HSV) activity and treatment for muco-cutaneous HSV infection (Yoosook et al. 2000).

AP is a common medicinal plant used in asian countries to treat various diseases like pharyngitis, tonsillitis, upper respiratory tract infection and acute monocytic leukemia (Thamlikitkul et al. 1991; Saxena et al. 2010; Tan et al. 2017). Preparations containing AP extract reported to prevent and improve the symptoms of common cold (Saxena et al. 2010). The therapeutic effect of AP in common cold management has been demonstrated by several clinical trials over the past 30 years (Thamlikitkul et al. 1991; Melchior et al. 1997; Caceres et al. 1997; Melchior et al. 2000). The meta-analysis studies have also endorsed the same (Poolsup et al. 2004).

The compounds inophyllum B, inophyllum P isolated from CI were found to have anti-HIV-1 reverse transcriptase activity (Laure et al, 2008). The compound calanolide A from CI has also been reported to inhibit HIV-1 replication (Kashman et al. 1992).

## 2. Materials and Methods

### 2.1. Virtual screening of plant molecules against SARS-CoV-2 targets

#### 2.1.1. Ligand preparation

Considering the availability and sustainability of medicinal plants, 521 Indian medicinal plants were identified and the structure of phytochemicals from each plant were manually curated from a reported research article of high quality. The biological active conformations of 13,105 curated plant molecules were obtained through ligand preparation process. The structure of plant molecules were sketched using a 2D sketcher tool available in Schrodinger maestro and subjected for ligand minimisation using Ligprep (LigPrep, version 2.3, Schrödinger, LLC, New York, 2009). During the minimisation, force field OPLS_2005 were assigned and stereoisomers were calculated after retaining specific chiralities.

#### 2.1.2. Protein Preparation

The proteins involved in, entry, replication and assembly of SARS-CoV-2 in human host such as Spike glycoprotein (S-Protein), RNA dependent RNA polymerase (RdRp), Main protease (Mpro), and 3-chymotrypsin like protease (3CLpro) were identified as a potential drug targets (Bharath et al. 2020; Yin et al. 2020; Jin et al. 2020; Douangamath et al. 2020). The structure of S-Protein was modelled as reported in Bharath et al. 2020 and the structure of remaining target proteins were downloaded from Protein Data Bank (PDB). The details of PDB ID are provided in **Table 1**. The three-dimensional structure of target proteins were prepared using the protein preparation wizard workflow available in the Schrödinger 2019-2 glide module. During the protein preparation, the crystallographic water molecules with less than three H-bonds were deleted and hydrogen atoms corresponding to neutral pH were added in consideration of ionisation states of amino acids. Following this, coordinates for any missing side-chain atoms were added using Prime v4.0, Schrödinger 2019-2 and energy minimisation was performed using the OPLS_2005 force field.

**Table 1:**
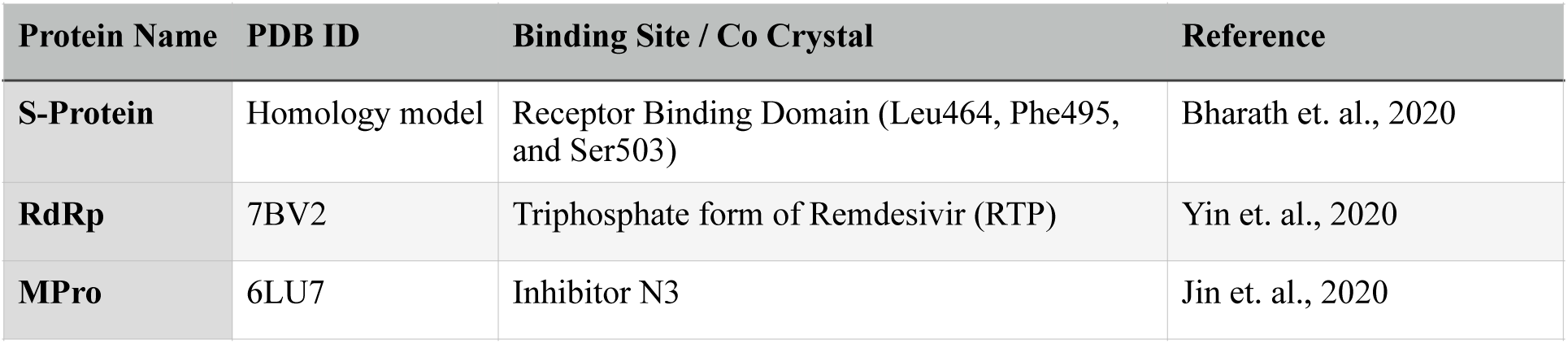
The details on structure of SARS-CoV-2 target proteins

#### 2.1.3. Virtual screening

The active site on the prepared receptor was confined with a 10Å radius by centering around selected residues for S-Protein and centering co-crystal molecules for other three target proteins. This generated a grid box measuring 20X20X20Å. The docking of phytochemicals over target proteins was performed using Glide v7.8. Schrödinger 2019-2 in different modes sequentially with defined and incremental precision, and computational time differences. The phytochemicals with best docked conformer and minimum glide energy were shortlisted for the evaluation of *in vitro* Anti-SARS-CoV-2 activity.

### 2.2. Extraction, Isolation and purification of plant molecules

#### 2.2.1. Plant materials, chemicals and reagents

The aerial parts of *ZP*, *CI, AP* and *CA,* and root part of *WS* were procured from Herbo Ayurvedics Calicut Kerala and authenticated by using herbarium, maintained in our laboratory. Organic solvents such as Hexane, EtOAc, DCM, MeOH, and Silica gel (60-120 mesh) for sample preparation were procured by Avra Synthesis Pvt Ltd. Silica high performance Gold columns were purchased from Redi sepRf, Teledyne ISCO. Solvents for HPLC, HPLC-grade Acetonitrile, Water and MeOH were procured from Sigma-Aldrich (Merck).

#### 2.2.2. Extraction and primary analysis of crude extracts

The plant materials were subjected for extraction using conventional bio-separation techniques.The 500g of plant material was packed in Soxhlet extractor, defatted by using hexane for about four hours. The dried defatted plant material was again subjected for extraction using 100% Methanol. The extraction was performed until the solvent in the thimble became colourless. The total filtrate was then concentrated by rotary evaporation under vacuum to obtain the ethanol crude extracts and the yield of the crude extracts was measured (**Table 2**). The 0.1g of crude extracts were dissolved in 10ml of methanol and subjected for thin layer chromatography using silica gel as stationary phase and different sets of solvent system as mobile phase (**Table 3**).

**Table 2:**
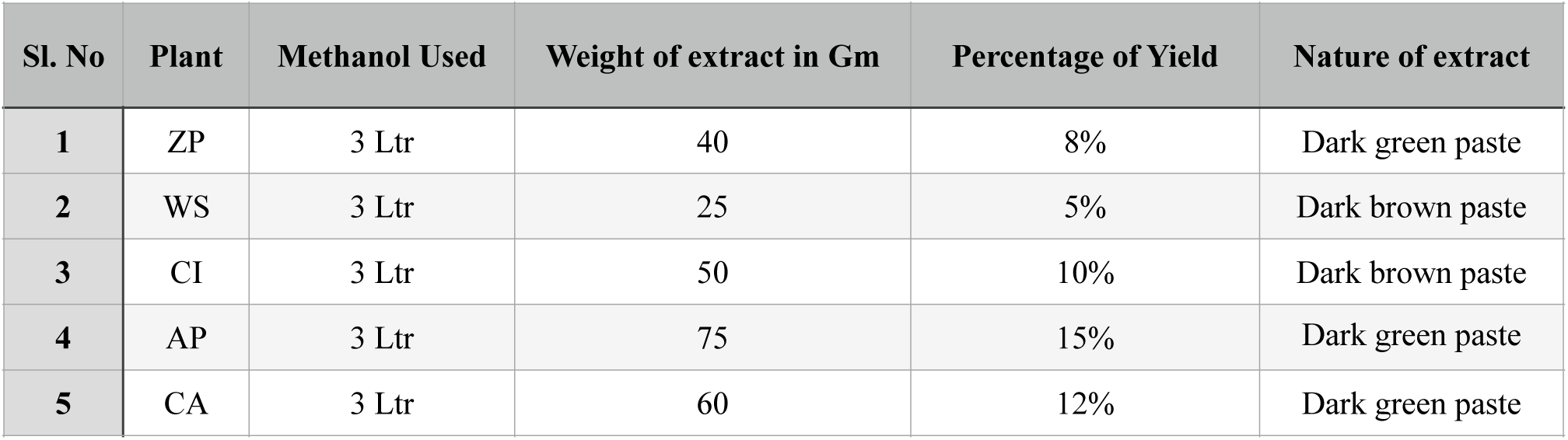
The details on yield and nature of crude methanol extracts.

| Sl. No | Plant | Methanol Used | Weight of extract in Gm | Percentage of Yield | Nature of extract |
| --- | --- | --- | --- | --- | --- |
| 1 | ZP | 3 Ltr | 40 | 8% | Dark green paste |
| 2 | WS | 3 Ltr | 25 | 5% | Dark brown paste |
| 3 | CI | 3 Ltr | 50 | 10% | Dark brown paste |
| 4 | AP | 3 Ltr | 75 | 15% | Dark green paste |
| 5 | CA | 3 Ltr | 60 | 12% | Dark green paste |

**Table 3:** The details on solvent system used and outcomes of thin layer chromatography analysis performed for crude methanol extracts.

| Sl. No | Plant | Solvent system | Proportion | Spot Number | Rf | Tota distance in cm |
| --- | --- | --- | --- | --- | --- | --- |
| 1 | ZP | Ethyl acetate:MeOH | 90:10 | 1 | 0.65 | 4.0 |
|  |  |  |  | 2 | 0.92 |  |
| 2 | WS | DCM:MeOH | 95:05 | 1 | 0.50 | 4.8 |
|  |  |  |  | 2 | 0.60 |  |
|  |  |  |  | 3 | 0.66 |  |
| 3 | CI | DCM:MeOH | 95:05 | 1 | 0.63 | 4.4 |
|  |  |  |  | 2 | 0.77 |  |
|  |  |  |  | 3 | 0.95 |  |
| 4 | AP | Ethyl acetate:MeOH | 80:10 | 1 | 0.32 | 5.00 |
|  |  |  |  | 2 | 0.40 |  |
| 5 | CA | Ethyl acetate:MeOH | 95:05 | 1 | 0.30 | 5.3 |
|  |  |  |  | 2 | 0.39 |  |
|  |  |  |  | 3 | 0.79 |  |

#### 2.2.3. Liquid chromatography–mass spectrometry (LC-MS) based separation and purification

The crude extracts were subjected for flash chromatography (Teledyne ISCO, CombiFlash NEXTGEN 300 system) using Redisep C-18 86G column for semi purification. Acetonitrile, water, and methanol were the solvents used at different concentrations volume by volume. The semi purified fractions were further subjected to HPLC for next level purification. The HPLC system used was WatersTM e2695 consisting of a quaternary pump, an automatic degasser, and an auto-sampler, having PDA and UV detector, column used was Shimpack C-18 (ODS) (250mm*4.6,5µm) column. The molecular weight of purified fractions were identified using the Shimadzu2020 LC-MS system. The temperature of the column was set to 40 °C and a 5 µL aliquot of the sample solution was injected at specific flow rate vis., 0.3ml/min,1ml/min, 1ml/min, 1ml/min, and 0.3ml/min for ZP, WS, CI, AP and CA fractions respectively. The gradient system followed for each sample was 70:30 for Water: Acetonitrile, 100% for Acetonitrile, and 70:30 for 0.1% formic acid in Water: 0.1% formic acid in MeOH. The positive ionization mode was used for compound ionization. The quantification was obtained in multiple reaction monitoring (MRM) mode with the precursor-product ion transition. High-purity nitrogen (N2) was used as the nebulizing gas, and nitrogen (N2) was used as the drying gas at a flow rate of 15 L/min. The mass spectrometer was operated at a capillary voltage of 4000 V, source temperature of 100 °C and desolvation temperature of 350 °C.

Before sending the pure fractions for In-vitro studies, they were allotted code names: *Z. piperitum* as ATRI-CoV-E1, *W. somnifera* as ATRI-CoV-E2, *C. inophyllum* as ATRI-CoV-E3), *A. paniculata* as ATRI-CoV-E4), *C. Asiatica* as ATRI-CoV-E5. Then they were sent to Regional Centre for Biotechnology, Faridabad (RCBF) for in *vitro* studies.

### 2.3. *In vitro* cytotoxicity and Anti-SARS-CoV-2 activity

In vitro cytotoxicity and Anti-SARS-CoV-2 activity was carried out at RCBF, using the methods described below:

#### 2.3.1. Preparation of cells

The cytotoxicity assay was done in a 96-well plate, with 3 wells for each sample. 1x10e4 VeroE6 cells were plated per well and incubated at 37-degree C overnight for the monolayer formation. Next day, cells were incubated with the test substance (TS) at the indicated concentration (**Table 4**) in 0.5% Dimethyl sulfoxide (DMSO). The control cells were incubated without TS.

**Table 4:** The dosage of plant extracts evaluated for cytotoxicity and anti-ATRI-CoV-2 activity.

| TS | Plant Source | Concentration (μM). |
| --- | --- | --- |
| Remdesevir | Synthetic | 1 μl added from 10 mg/ml solution in 200 μl |
| ATRI-CoV-E1 | Z. piperitum | 1 μl added from 30 mg/ml solution in 200 μl |
| ATRI-CoV-E2 | W. somnifera | 1 μl added from 2.8 mg/ml solution in 200 μl |
| ATRI-CoV-E3 | C. inophyllum | 1 μl added from 5 mg/ml solution in 200 μl |
| ATRI-CoV-E4 | A. paniculata | 1 μl added from 5.5 mg/ml solution in 200 μl |
| ATRI-CoV-E5 | C. asiatica | 1 μl added from 45 mg/ml solution in 200 μl |

#### 2.3.2. In vitro cytotoxicity

The treated incubated cells were collected at 24 and 48 hours and stained with Hoechst 33342 and Sytox orange dye. 16 images per well were taken at 10X, which covered 90% of well area using ImageXpress Microconfocal (Molecular Devices). The percentage of cell viability was measured by counting the number of dead cells in the Sytox image as described by Tan et al. 2004.

#### 2.3.3. In vitro Anti-SARS-CoV-2 activity

The prepared cells were infected with SARS-CoV2 at a multiplicity of infection (MOI) of 0.01. After 24 and 48 hours, viral RNA was extracted from 100 µl culture supernatant and subjected to qRT-PCR where Ct values for N and E gene sequence were determined. Inhibition of virus replication was determined by the Ct value in TS-treated cells compared to the control. Remdesivir was used as a positive control for viral inhibition (Caly et al. 2020).

#### 2.3.4. *In vitro* IC_50_ determination

The prepared cells were incubated with TS at seven different concentrations as follows. The concentrations used for ATRI-CoV-E4, were 5.5, 2.75, 1.37, 0.68, 0.34, 0.17 and 0.08 µg. The concentrations used for Remdesivir were 10, 3, 1, 0.3, 0.1, 0.03 and 0.01 µM. The control cells were incubated with 0.5% DMSO.

The cells were infected with SARS-CoV2 at a MOI of 0.01 and after 48 hours, viral RNA was extracted from 100 µl culture supernatant and subjected to qRT-PCR. The Ct values for N and E gene sequence were recorded and the inhibition of virus replication was determined based on the change in the Ct value in TS-treated cells compared to the control. IC_50_ values were determined using AAT Bioquest IC_50_ calculator.

### 2.4. *In vivo* acute and subacute toxicity

#### 2.4.1. Experimental animals

Toxicity assays were conducted in accordance with the standard guidelines of the Organization for Economic Cooperation and Development (OECD, 2001) for use of animals in scientific research. Experimental animals were housed in groups of three for acute toxicity study and a group of six animals for sub-acute toxicity study as per standard laboratory conditions. Detail of animals is given in **Table 5**.

**Table 5:**
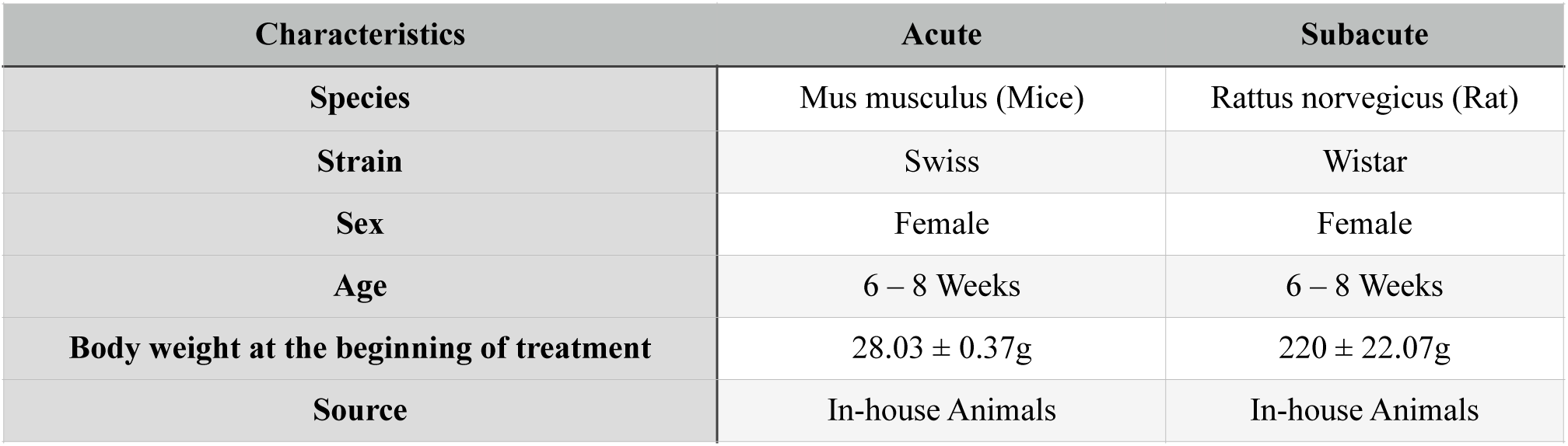
Detailed characteristics of animals used for acute and subacute toxicity studies.

| Characteristics | Acute | Subacute |
| --- | --- | --- |
| Species | Mus musculus (Mice) | Rattus norvegicus (Rat) |
| Strain | Swiss | Wistar |
| Sex | Female | Female |
| Age | 6 – 8 Weeks | 6 – 8 Weeks |
| Body weight at the beginning of treatment | 28.03 ± 0.37g | 220 ± 22.07g |
| Source | In-house Animals | In-house Animals |

The experimental female rats were bred and reared at the animal care facility in NUCARE, Nitte University, Mangalore and experiment was carried out in the same facility. Adult albino Wistar non-pregnant female rats weighing 200−220 g and between 8–10 weeks of age were used.

#### 2.4.2. Preparation of Dose Formulation

The test substance was prepared by standard procedure. Extracts weighing 833.33mg ATRI-CoV-E2, ATRI-CoV-E4 and ATRI-CoV-E5 each were placed in mortar. Small quantity of water was added, and the sample was triturated using a pestle. Once all samples were wet, remaining quantity of the water was added slowly to make up the total volume to 5ml, while trituration continued to get simple suspension and the dose formula was labeled as ATRICOV 452.

#### 2.4.3. Acute toxicity study

Three male rats and three female mice were selected and fasted overnight (12 h). Animals were not given food but had free access to water. The single dose of ATRICOV 452 at 2000 mg/kg was administered using an oral gavage needle. Animals were observed every 30 minutes for the first 4 hours, and daily thereafter, for a total of 14 days for toxic manifestations by observing clinical signs (salivation, excitability, draping, tremors, twitching, rising fur) and death. The weight and food consumption of the experimental animals were recorded on the first day and 14th day. After 14 days, they were sacrificed.

#### 2.4.4. Subacute toxicity study

As per OECD (2001) recommendations, the dose at which the extract is not expected to produce mortality or severe acute toxicity is called the starting dose of the sub-acute toxicity (Tilahun et al. 2020).

The 2000 mg/kg body weight of the ATRICOV 452 was the highest dose determined. A total of 72 Wistar male and female rats were divided into 6 groups as shown in **Table 6**; each group consisted of 6 male and 6 female rats. The groups five and six were called satellite groups. Each group was given 1 mL/100 g of body weight of ATRICOV 452 at concentrations of 0.00 mg/kg (control), 200 mg/kg (low dose), 400 mg/kg (medium dose), 800mg/kg (high dose), 0.00 mg/kg (control reversal) and 800mg/kg (high dose reversal). All the six groups received the treatment for 28 days. The weight and food consumption of animals were measured weekly till the 28th day. On day 29, control, low, medium and high dose groups were sacrificed. The satellite groups were maintained for another 14 days without any treatment to observe the reversible untoward effect of treatment if any. On day 43, satellite groups were sacrificed for hematological, biochemical and histopathological examinations.

**Table 6:** Grouping of animals and dose for sub-acute toxicity studies

| Group | Treatment Group | Dose (mg/kg) | Dose Conc. (mg/ml) | Animals/Group |  | Total No. of Animals/Group |
| --- | --- | --- | --- | --- | --- | --- |
|  |  |  |  | Male | Female |  |
| **G I** | Control | 0 | --- | 6 | 6 | 12 |
| **G II** | Low | 200 | 50 mg/ml | 6 | 6 | 12 |
| **G III** | Mid | 400 | 100 mg/ml | 6 | 6 | 12 |
| **G IV** | High | 800 | 200 mg/ml | 6 | 6 | 12 |
| **G V** | Control - Rev | 0 | --- | 6 | 6 | 12 |
| **G VI** | High - Rev | 800 | 200 mg/ml | 6 | 6 | 12 |

##### 2.4.4.a. Haematological studies

1 mL of blood samples were collected from each rat. The hematological examination was carried out to measure differences in Red Blood Cell (RBC), Packed cell volume (PCV), Haemoglobin, Neutrophils, Basophils, Lymphocytes and Monocytes counts.

##### 2.4.4.b. Biochemical markers

The levels of glucose, Aspartate aminotransferase (AST), Alanine aminotransferase (ALT), Alkaline phosphatase (ALP), Total Protein (TP), Albumin, Total cholesterol (T.Chol), Triglyceride (TG), Blood urea nitrogen (BUN), creatinine, total bilirubin, calcium, phosphate, sodium and potassium were measured.

##### 2.4.4.c. Histopathological examinations

All rats were sacrificed using Isoflurane anesthesia at the end of the 14th day for acute toxicity and at the end of the 28th day for sub-acute toxicity studies. Macroscopic and histopathological examinations were done for liver, kidney and jejunum.

#### 2.4.5. Statistical analysis

The data were analysed using Graph Pad Prism version 6 Software. For each group, one way ANOVA analysis was done followed by Dunnett’s comparison test. The data were expressed as mean ± standard deviation (SD). A p-value < 0.05 was considered as statistically significant.

## 3. Results and discussion

### 3. 1. Virtual Screening

The virtual screening of 13,105 phytochemicals at different precision levels like high throughput virtual screening (HTVS), standard precision (SP) and extra precision (XP) has suggested the interaction of phytochemicals from ZP, WS, CI, AP and CA with SARS-CoV-2 drug targets.

The SARS-CoV-2 enters the host by interacting its S-Protein with a host receptor called ACE2. The S-protein consists of two subunits, S1 as the receptor-binding domain (RBD) and S2 subunit as the fusion protein. The residues forming RBD of SARS-CoV-2 Leu455, Phe486, and Ser494 (corresponding to Leu464, Phe495, and Ser503 of S-protein of SARS-CoV-2 modelled in our earlier study (Bharath et al. 2020)) are and known to play a critical role in disease transmission (Wan et al. 2020). SARS-CoV-2 recognises human ACE2 by its residues Gln493 and Leu455 (corresponds to Gln502 and Leu464), thereby enhancing viral binding to human ACE2. Hence, the interaction of phytochemicals with Gln502 or Leu464 can inhibit the interaction of S-Protein with the ACE2 receptor and stop the entry of SARS-CoV-2 into the host cell. The molecular docking has shown the interaction of the physicochemical called Kaempferol from CA with the residues Gln502 and Val492 making the ridges which are sterically complementary to the cleft on the human ACE2 receptor (**Figure 1A**). The H-bonds formed between hydroxyl groups of Kaempferol and Nγ of Gln502 side chain and ketone group of Val492 backbone residues showed strong interaction (**Figure 1B**). The major contribution for docking energy was contributed by H-Bonds (**Table 7**) and the penalty was due to high solvent exposure of Kaempferol on the surface of S-Protein (**Figure 1C**).

**Figure 1:**
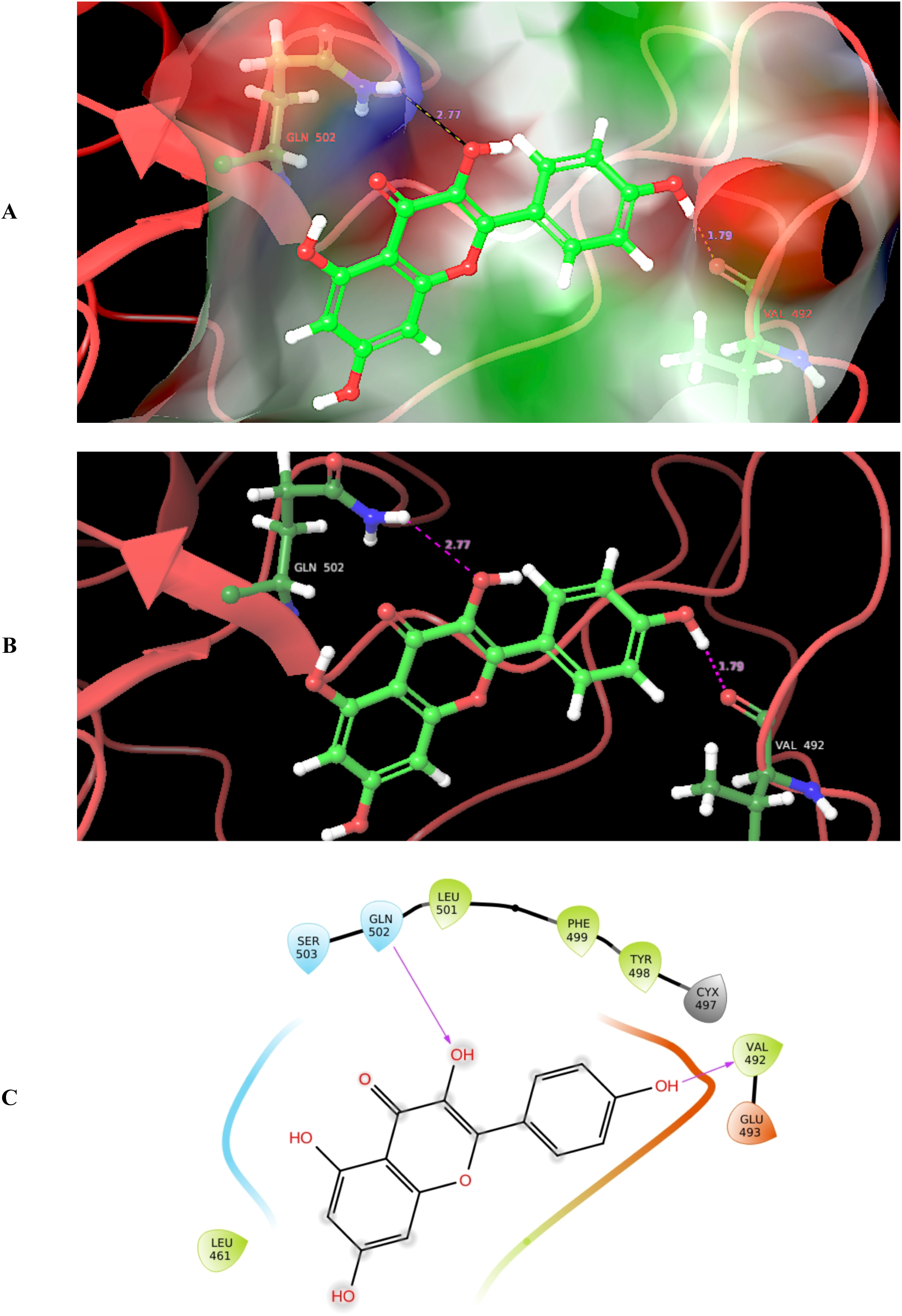
Molecular docking of Kaempferol (Green coloured ball and stick) in the RBD site of SARS-CoV-2 S-Protein. **A:** Three dimensional view where receptor is rendered with molecular surface to illustrate the interaction of Kaempferol with the amino acids Gln502 and Val492 making the ridges which are sterically complementary to the cleft on human ACE2 receptor. **B: 3**D view illustrating the interaction of Kaempferol hydroxyl groups with Nγ of Gln502 side chain and ketone group of Val492 back bone residues. **C:** The protein ligand interaction diagram illustrating the interaction of Kaempferol in the RBD site of SARS-CoV-2 S-Protein; the grey highlights on ligand molecules represents solvent exposure.

**Table 7:** List of hits generated by molecular docking of phytochemicals against three major SARS-CoV-2 targets

| Phytochemical | Plant Source | Target | No. of H-Bonds | H-Bond forming Atoms | | Bond length in Å | Docking Energy $\text{mJ/mol}$ |
| --- | --- | --- | --- | --- | --- | --- | --- |
|  |  |  |  | Ligand | Target |  |  |
| Kaempferol | CA | S-Protein | 2 | C4': OH | Val492: O | 1.79 | -5.21 |
| | | | | C7: OH | Gln502: Nγ | 2.77 | |
| Andrographidine C | AP | RdRp | 5 | C2'': OH | U20: C3:OH | 2.49 | -8.12 |
|  |  |  |  | C4: O | U20: C3:OH | 2.16 |  |
| | | | | C6'': OH | Lys551: Hζ1 | 2.33 | |
| | | | | C4'': OH | Glu811: Oε2 | 2.47 | |
|  |  |  |  | C6'': OH | POP 1003: O2 | 2.63 |  |
| Coagulin G | WS | MPro | 3 | C17: OH | Thr26: O | 2.04 | -9.17 |
| | | | | C3: OH | Ser144: Hγ | 2.39 | |
| | | | | C1: O | His163: Nε2 | 2.55 | |
| Zanthoxyl flavone | ZP | MPro | 4 | C5: OH | Thr26: O | 1.96 | 8.23 |
|  |  |  |  | O | Gly143: N | 3.71 |  |
| | | | | C4': OH | His163: Nε2 | 2.02 | |
| | | | | C3: OH | Gln189: Oε1 | 2.73 | |
| 3,4,6,8-Tetrahydroxy-7-(3-hydroxy-3-methylbutyl)-9H-xanthene-9-one | CI | MPro | 3 | C11:OH | Thr25: Oγ1 | 1.81 | 8.75 |
|  |  |  |  | C6: OH | Thr26: O | 2.28 |  |
| | | | | C3: OH | His163: Nε2 | 1.80 | |

The inhibition of SARS-CoV-2 replication in cell-based assays are demonstrated for small molecule antiviral nucleoside analog drugs like Remdesivir, Favipiravir, Ribavirin, and Galidesivir (Lu et al. 2020; Sheahan et al. al., 2020. The nucleotide analog drugs are reported to inhibit the viral viral enzyme called RdRp through non-obligate RNA chain termination mechanism, and the crystal structure of the template-RTP RdRp complex provides an experimental model to rationalise how these drugs inhibit the SARS-CoV-2 RdRp activity. In the present study, the conformation of the Andrographidine C from AP has also shown non-obligate RNA chain termination type of interaction (**Figure 2)** and the interaction was similar to the co-crystal (**Figure 2D**) . The orientation of Andrographidine C was parallel to the base Uracil 20 (U 20) of RNA and the interaction was found stable due to pi-pi stacking between coumarin moiety and Uracil base. Additionally, the H-Bonds formed by electron donating hydroxyl groups present in the Andrographidine C have strengthened the interaction with minimum docking energy (**Table 7**).

**Figure 2:**
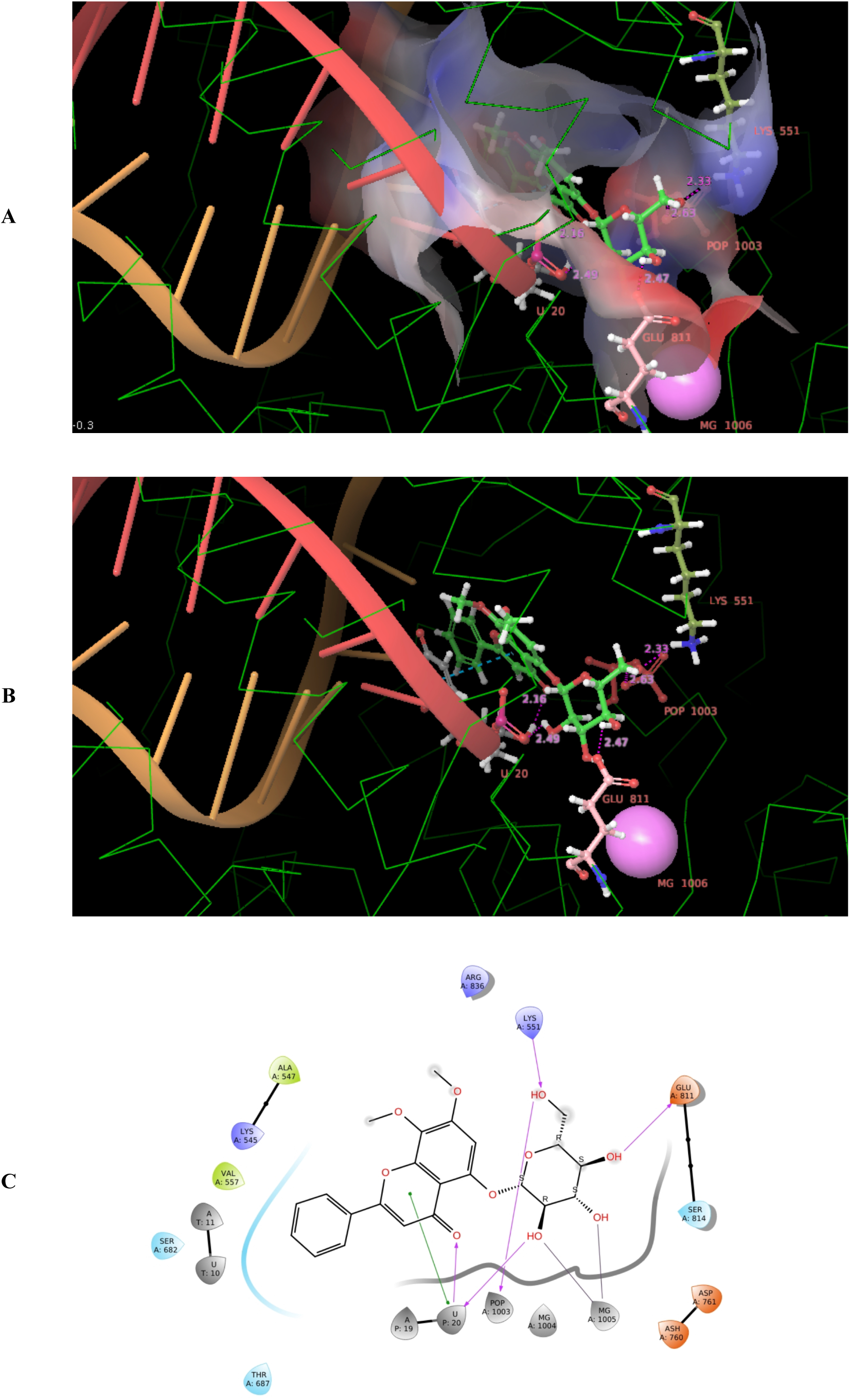
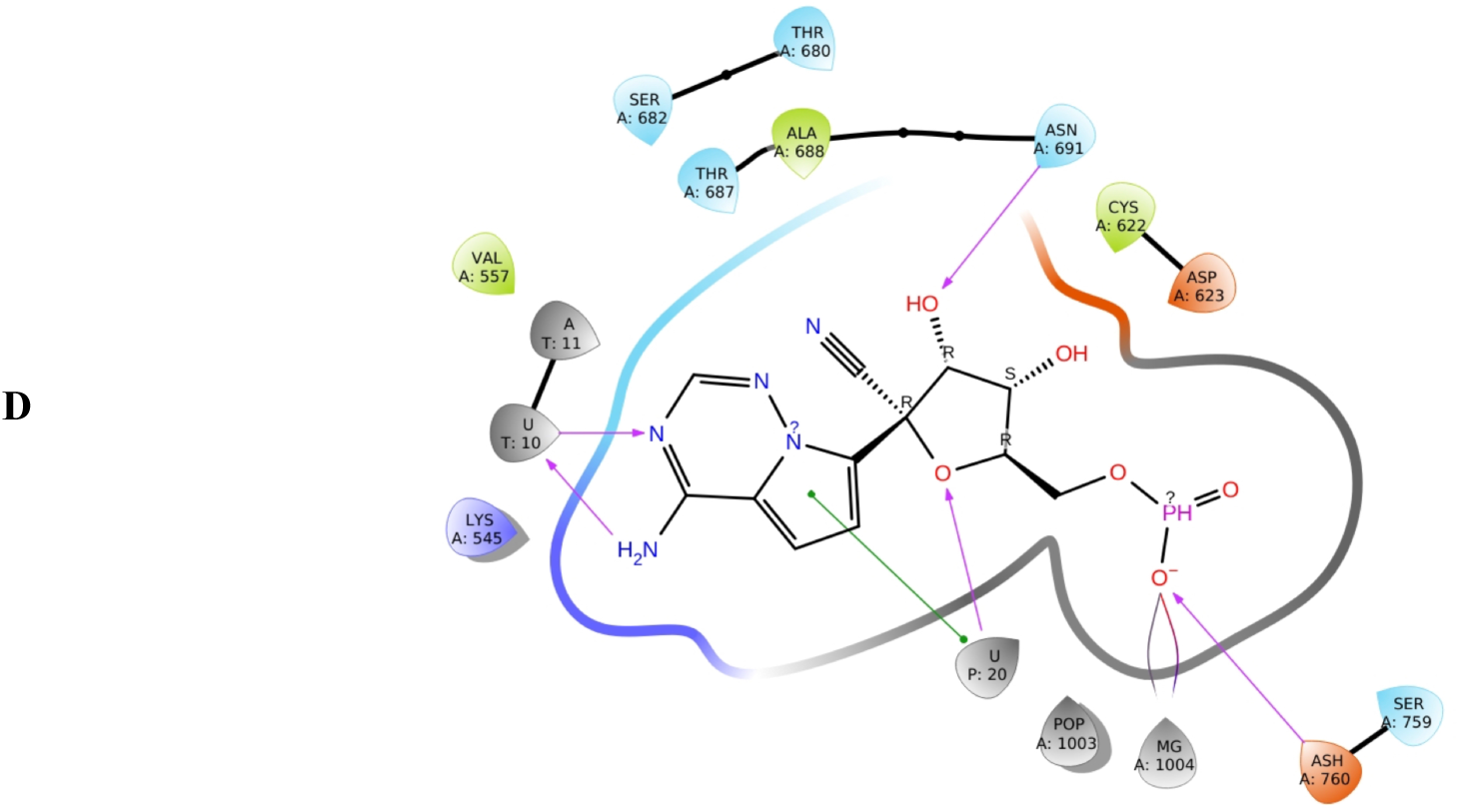
Molecular docking of Andrographidine C (Green coloured ball and stick)in the RNA binding site of SARS-CoV-2 RdRp **A:** Three dimensional view where RDRP RNA binding site is rendered with molecular surface to illustrate the interaction of Andrographidine C with the amino acids Lys551and Glu811; non-obligate RNA chain termination due to interaction with Uracil 20 is also notable. **B: 3**D view illustrating the H-bonds between hydroxyl groups of Andrographidine C and ribose group of U20, Oε2 of Glu811 and Hζ1 of Lys551; the pi-pi stacking (blue dotted line) between coumarin group of Andrographidine C and Uracil 20 is notable. **C:** The protein ligand interaction diagram illustrating the interaction of Andrographidine C in the RNA binding site of SARS-CoV-2 RdRp. **D:** The protein ligand interaction diagram illustrating the interaction of the co-crystal RTP in the RNA binding site of SARS-CoV-2 RdRp; the grey highlights on ligand molecules represents solvent exposure.

Jin et al. have elucidated the inhibitory mechanism of N3 by determining the crystal structure of SARS-CoV-2 Mpro in complex with N3 (Jin et al. 2020). The elucidated crystal structure is a homodimer with two designated protomers A and B. The specific interactions of N3 with Mpro showed the covalent bond between Sγ atom of Cys145 and Cβ atom of the vinyl group in N3. The side chains of Phe140, Asn142, Glu166, His163 and His172 of protomer A, and the backbone atoms of Phe140 and Leu141 of protomer A are involved in the formation of the S1 subsite. The lactam at P1 of N3 inserts into the S1 subsite and forms a hydrogen bond with His163 of protomer A. Similarly, the Coagulin G from WS, Zanthoxyl flavone from ZP and 3,4,6,8-Tetrahydroxy-7-(3-hydroxy-3-methylbutyl)-9H-xanthen-9-one from CI showed interaction with His163 (**Figure 3, 4 and 5)**. The interactions were favoured by other residues in the cleft (**Table 7**).

**Figure 3:**
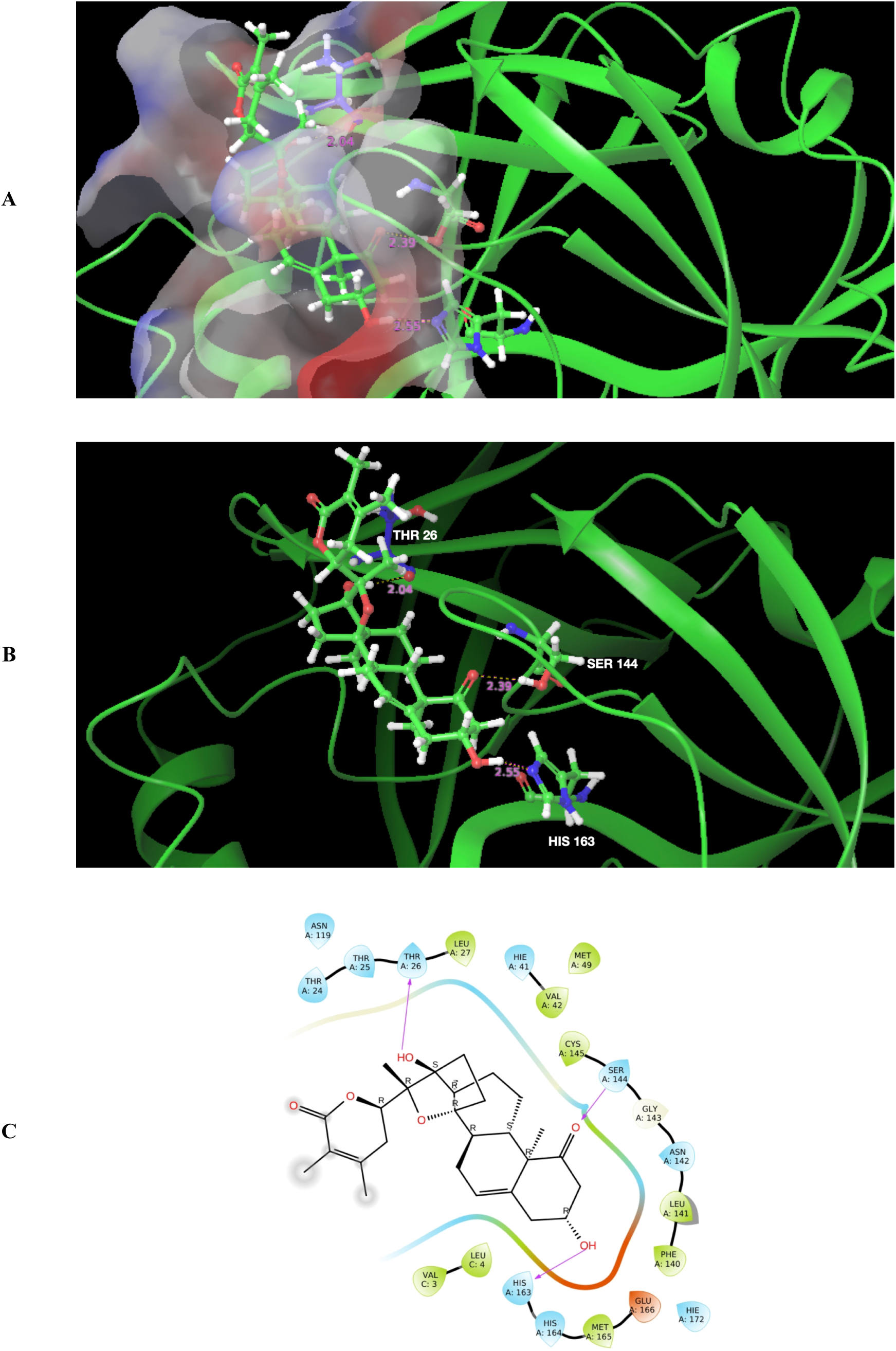
Molecular docking of Coagulin G from WS (Green coloured ball and stick) in the the S1 subsite of SARS-CoV-2 MPro. **A:** Three dimensional view where receptor is rendered with molecular surface to illustrate the interaction of Coagulin G in the the S1 subsite. **B: 3**D view illustrating the interaction of Coagulin G hydroxyl groups with Nε2 of His163, Hγ of Ser144 side chains and ketone back bone atom of Thr26 . **C:** The protein ligand interaction diagram illustrating the interaction of Coagulin G in the the S1 subsite of SARS-CoV-2 MPro; the grey highlights on ligand molecules represents solvent exposure.

**Figure 4:**
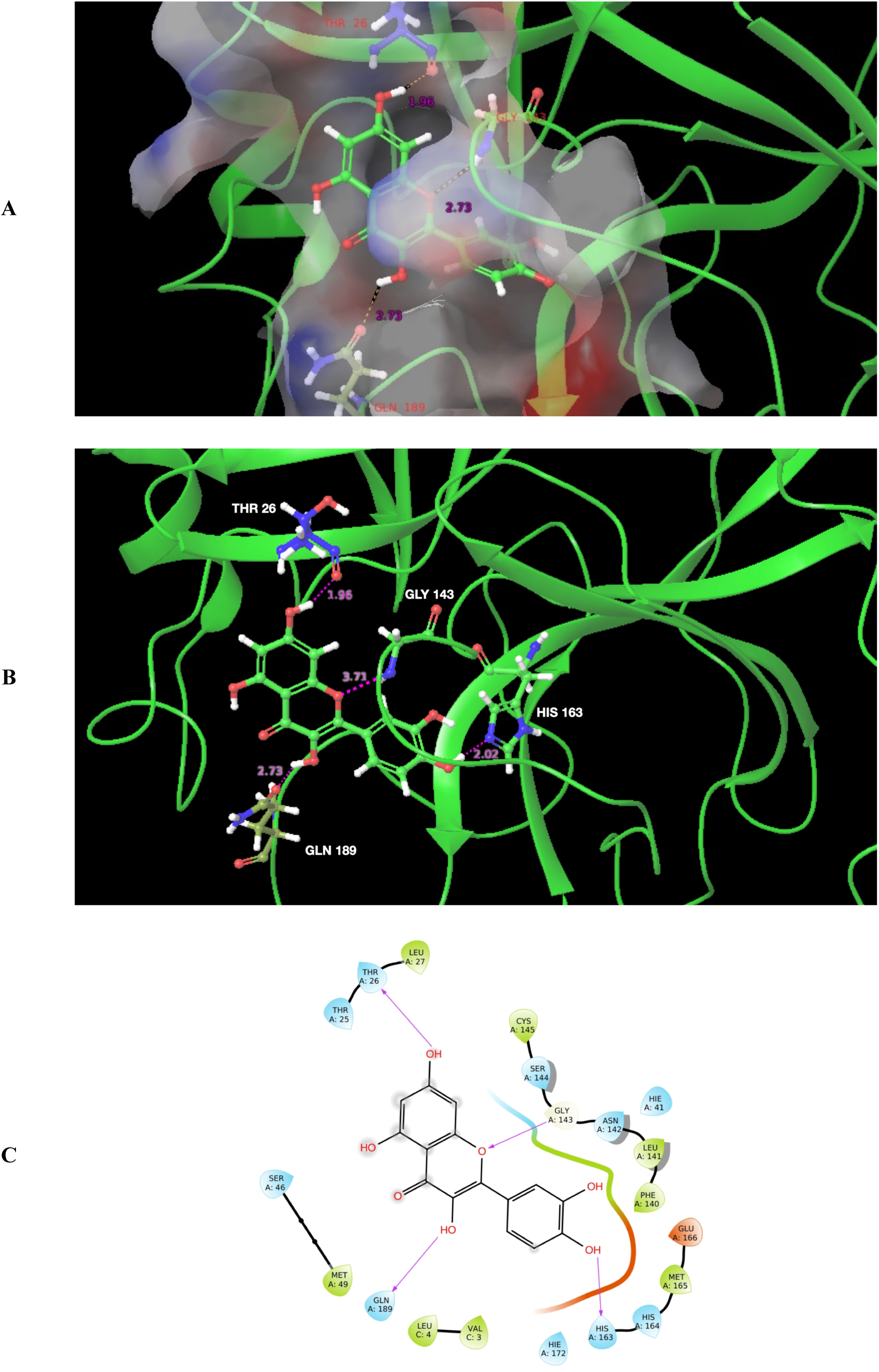
Molecular docking of Zanthoxyl flavone from ZP (Green coloured ball and stick) in the the S1 subsite of SARS-CoV-2 MPro. **A:** Three dimensional view where receptor is rendered with molecular surface to illustrate the interaction of Zanthoxyl flavone in the the S1 subsite. **B:** 3D view illustrating the H-Bond formed between Zanthoxyl flavone hydroxyl groups and Nε2 of His163, Oε1 of Gln189 side chains and ketone back bone atom of Thr26; O, the member of heterocyclic ring in the coumarin group of Zanthoxyl flavone formed the H-Bond with amine back bone atom of Gly143. **C:** The protein ligand interaction diagram illustrating the interaction of Zanthoxyl flavone in the the S1 subsite of SARS-CoV-2 MPro; the grey highlights on ligand molecules represents solvent exposure.

**Figure 5:**
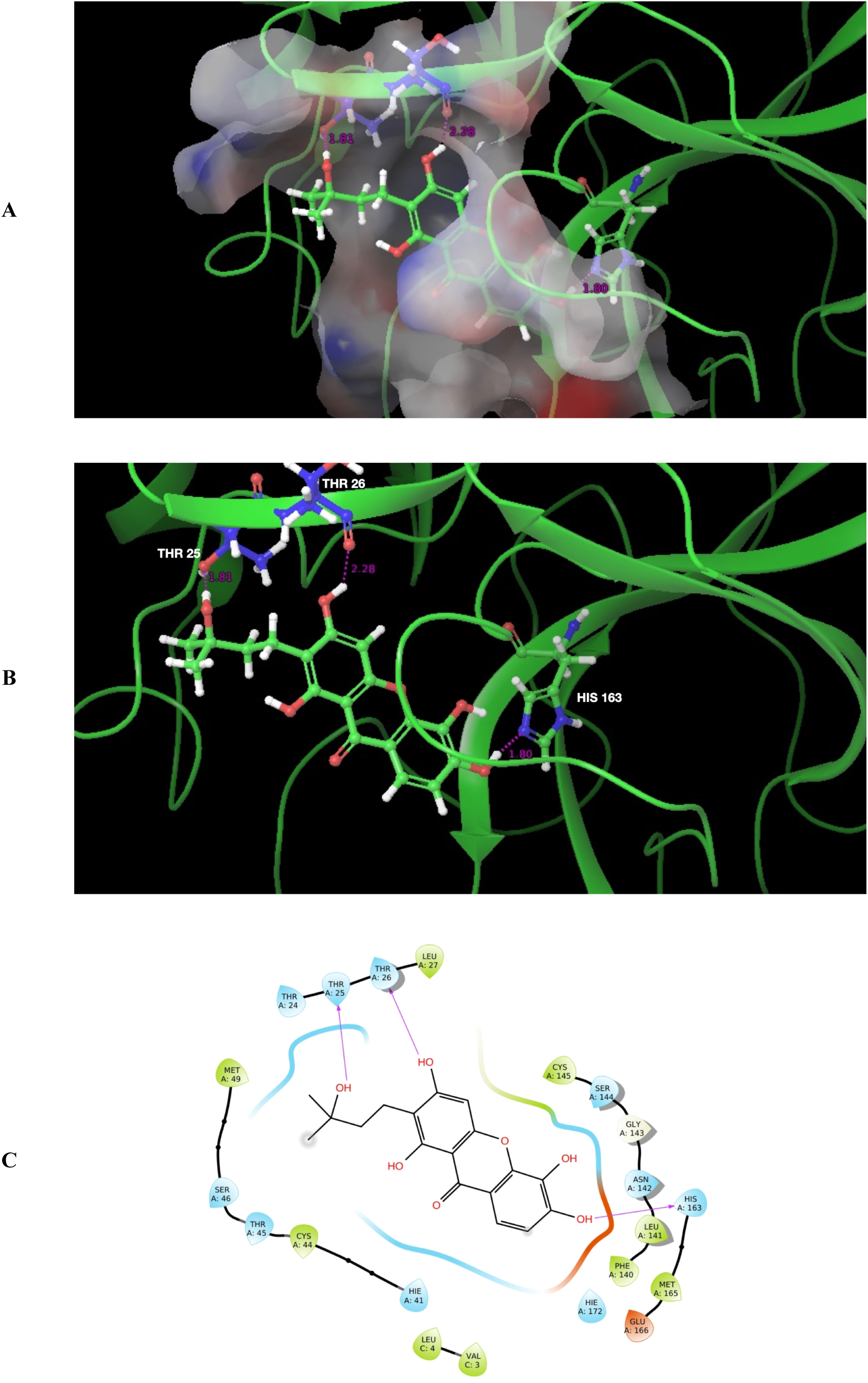
Molecular docking of 3,4,6,8-Tetrahydroxy-7-(3-hydroxy-3-methylbutyl)-9H-xanthene-9-one from CI (Green coloured ball and stick) in the the S1 subsite of SARS-CoV-2 MPro. **A:** Three dimensional view where receptor is rendered with molecular surface to illustrate the interaction of 3,4,6,8-Tetrahydroxy-7-(3-hydroxy-3-methylbutyl)-9H-xanthene-9-one in the the S1 subsite. **B:** 3D view illustrating the H-Bond formed between 3,4,6,8-Tetrahydroxy-7-(3-hydroxy-3-methylbutyl)-9H-xanthene-9-one hydroxyl groups and Nε2 of His163, Oγ1 of Thr25 side chains and ketone back bone atom of Thr26. **C:** The protein ligand interaction diagram illustrating the interaction of 3,4,6,8-Tetrahydroxy-7-(3-hydroxy-3-methylbutyl)-9H-xanthene-9-one in the the S1 subsite of SARS-CoV-2 MPro; the grey highlights on ligand molecules represents solvent exposure.

### 3.2. Extraction, Isolation and purification of plant molecules

The fractions of interest were purified using flash chromatography and confirmed for the presence of phytochemicals Zanthoxyl flavone, Coagulin G, 3,4,6,8-Tetrahydroxy-7-(3-hydroxy-3-methylbutyl)-9H-xanthen-9-one, Andrographidine C, and Kaempferol in ZP, WS, CI, AP and CA methanol extracts respectively.

The flash chromatographic fractionation of crude methanol extracts of ZP has led to four different polarity fractions (**Figure 6A**) at different RT (**Table 8**). The HPLC chromatogram for the first fraction obtained from flash chromatography at Rt1 of 25min has shown a high resolution peak at the retention time of 6.4 min based on UV wavelength at 210 nm (**Figure 6B**). The HPLC fraction obtained at the retention time of 6.4 min has shown a peak corresponding 347 m/z in MS spectra (**Figure 6C**) has ensured the presence of Zanthoxyl flavone in ZP methanol extract. Hence, it is inferred that, the Zanthoxyl flavone in the extract can interact with Mpro of SARS-CoV-2 and offer the Anti-SARS-CoV-2 activity.

**Figure 6:**
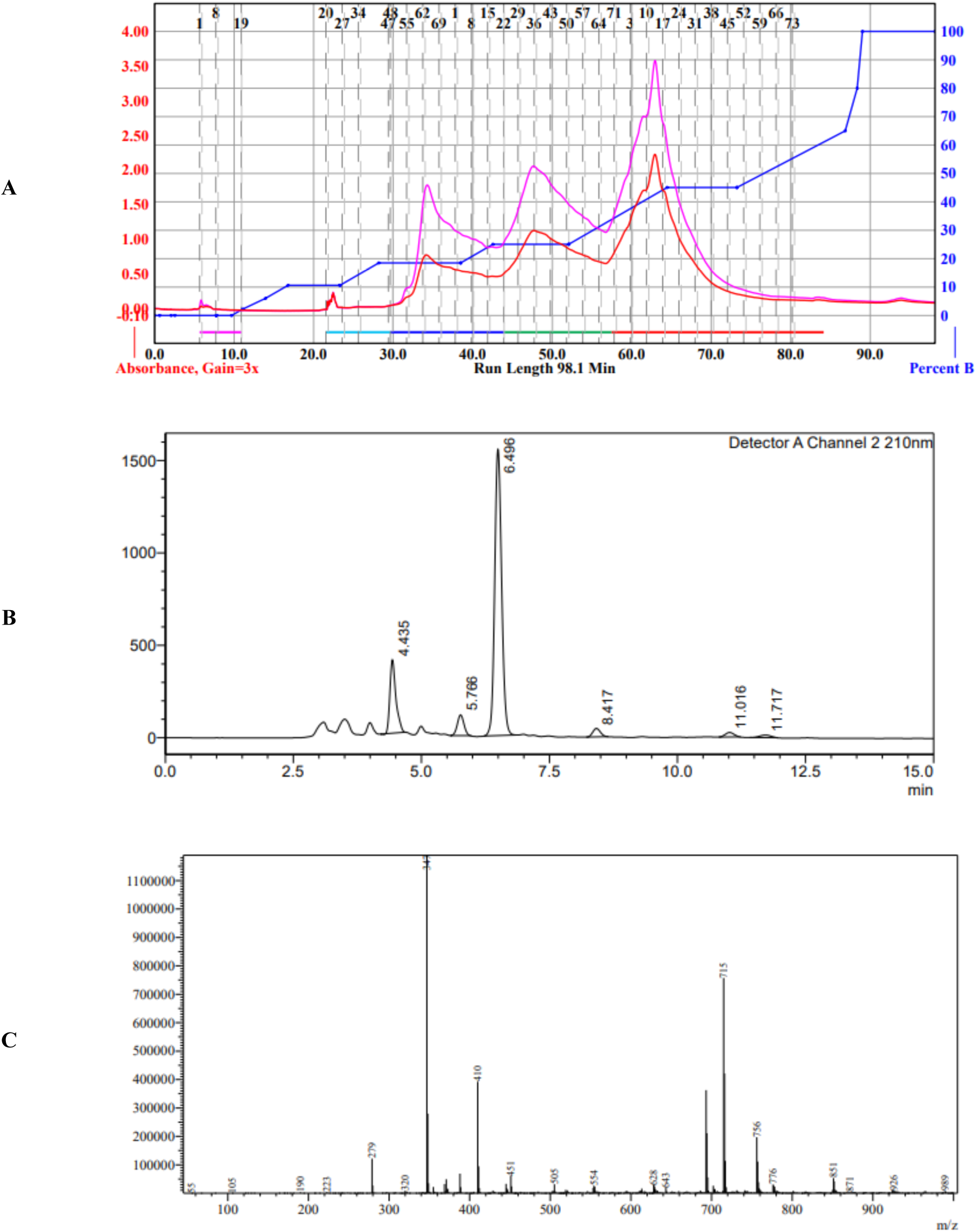
Chromatograms and spectra of ZP plant extracts, **A:** Flash chromatogram of the fractions in ZP methanol extract, **B:** HPLC chromatogram of the fractions in WS methanol extract, **C:** LC-MS spectra

**Table 8:** The details on solvent system used and outcomes of flash chromatography and HPLC analysis performed for methanol extracts.

| Flash Chromatography |  |  |  |  | HPLC |  |  |  |  |
| --- | --- | --- | --- | --- | --- | --- | --- | --- | --- |
| Sl. No | Plant | Mobile phase | Flow rate (ml/min) | RT (min) | Mobile phase | Flow rate (ml/min) | Wavelength (nm) | Time (min) | RT (min) |
| 1 | ZP | FMP1 | 30 | Rt1 – 25<br>Rt2 – 35<br>Rt3 – 48<br>Rt4 – 63 | HMP1 | 0.3 | 210 | 15 | 6.4 |
| 2 | WS | FMP1 | 30 | Rt1 – 22<br>Rt2 – 25<br>Rt3 – 45 | HMP2 | 1 | 254 | 10 | 5.2 |
| 3 | CI | FMP1 | 25 | Rt1 – 5<br>Rt2 – 40<br>Rt3 – 80 | HMP2 | 0.3 | 254 | 19 | 19.8 |
| 4 | AP | FMP1 | 25 | Rt1 – 5<br>Rt2 – 25 | HMP2 | 0.3 | 210 | 15 | 6.5 |
| 5 | CA | FMP2 | 30 | Rt1 – 20<br>Rt2 – 30<br>Rt3 – 90<br>Rt3 – 140 | HMP1 | 1 | 370 | 26 | 8.7 |
|  | **Flash Chromatography:** System = Combiflash NextGen 300, Column = RediSep C-18 86G, Solvent used for preparation of sample = 100% Acetonitrile. (Mobile Phase: FMP1 = Water:MeOH (0-100%), FMP2 = 0.1% FA in water & 0.1% FA in MeOH.<br>**HPLC:** System = Waters™ e2695, Column = Shimpack C-18 (ODS), Dimension = 250mm*4.6*5μm, Solvent used for preparation of sample = 100% Acetonitrile, Detector = PDA. (Mobile Phase: HMP1 = 0.1% FA in water & 0.1% FA in MeOH, HMP2 = Water:Acetonitrile (0-100%)) |  |  |  |  |  |  |  |  |

The flash chromatographic fractionation of crude methanol extracts of WS has led to three different polarity fractions (**Figure 7A**) at different RT (**Table 8**). The HPLC chromatogram for the first fraction obtained from flash chromatography at Rt1 of 22min has shown a high resolution peak at the retention time of 5.2min (**Figure 7B**) based on UV wavelength at 254 nm. The HPLC fraction obtained at the retention time of 5.42min has shown a peak corresponding 465 m/z in MS spectra (**Figure 7C**) has confirmed the presence of Coagulin G in WS methanol extract. Hence, it is inferred that, the Coagulin G in the extract can interact with Mpro of SARS-CoV-2 and offer the Anti-SARS-CoV-2 activity.

**Figure 7:**
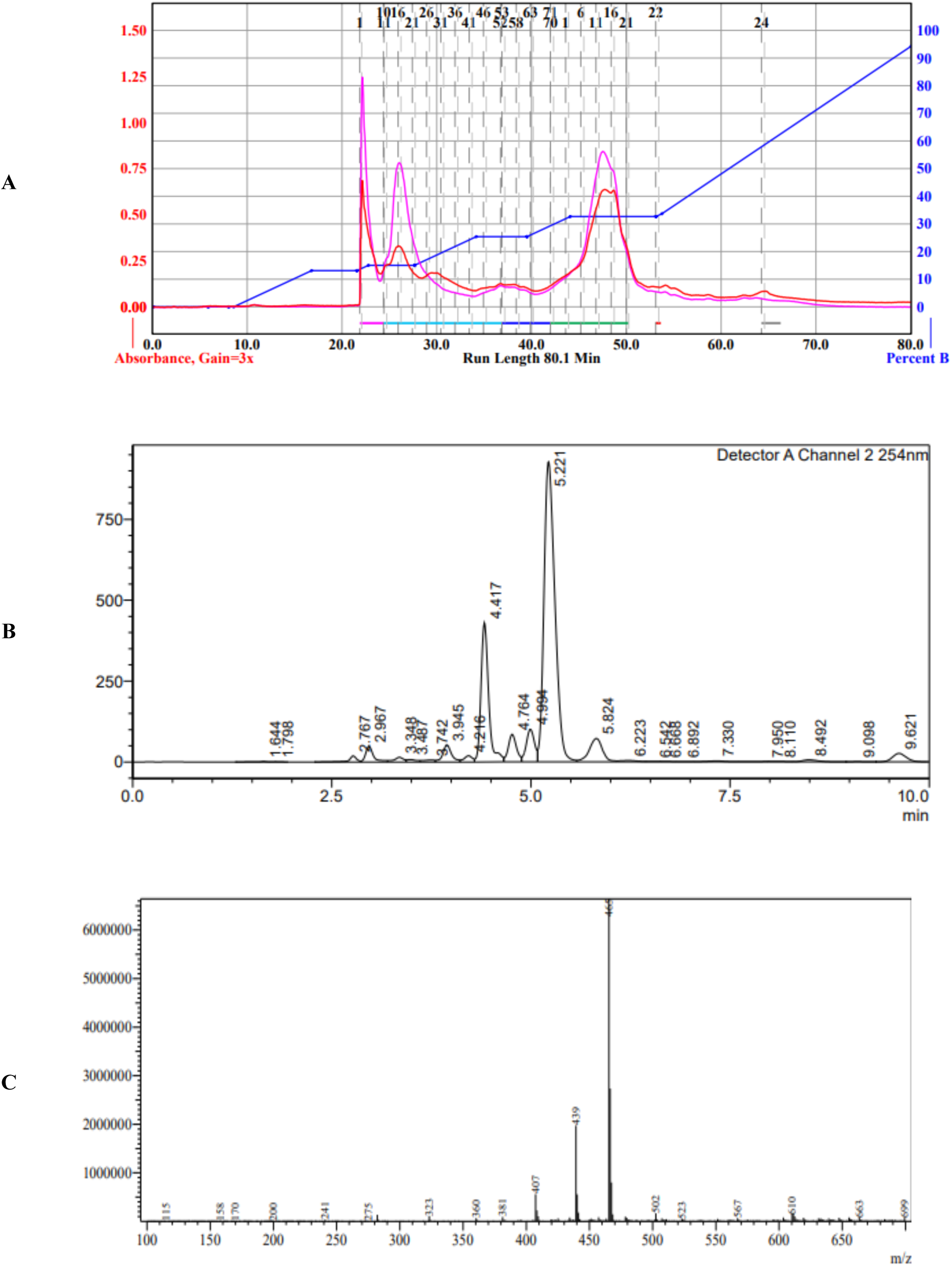
Chromatograms and spectra of WS plant extracts, **A:** Flash chromatogram of the fractions in WS methanol extract, **B:** HPLC chromatogram of the fractions in WS methanol extract, **C:** LC-MS spectra

The flash chromatographic fractionation of crude methanol extracts of CI has led to three different polarity fractions (**Figure 8A**) at different RT (**Table 8**). The HPLC chromatogram for the second fraction obtained from flash chromatography at Rt2 of 40min has shown a high resolution peak at the retention time of 19.8min (**Figure 8B**) based on UV wavelength at 254 nm. The HPLC fraction obtained at the retention time of 19.8min has shown a peak corresponding 344 m/z in MS spectra (**Figure 8C**) has confirmed the presence of 3,4,6,8-Tetrahydroxy-7-(3-hydroxy-3-methylbutyl)-9H-xanthene-9-one in CI methanol extract. Hence, it is inferred that, the 3,4,6,8-Tetrahydroxy-7-(3-hydroxy-3-methylbutyl)-9H-xanthen-9-one in the extract can interact with Mpro of SARS-CoV-2 and offer the Anti-SARS-CoV-2 activity.

**Figure 8:**
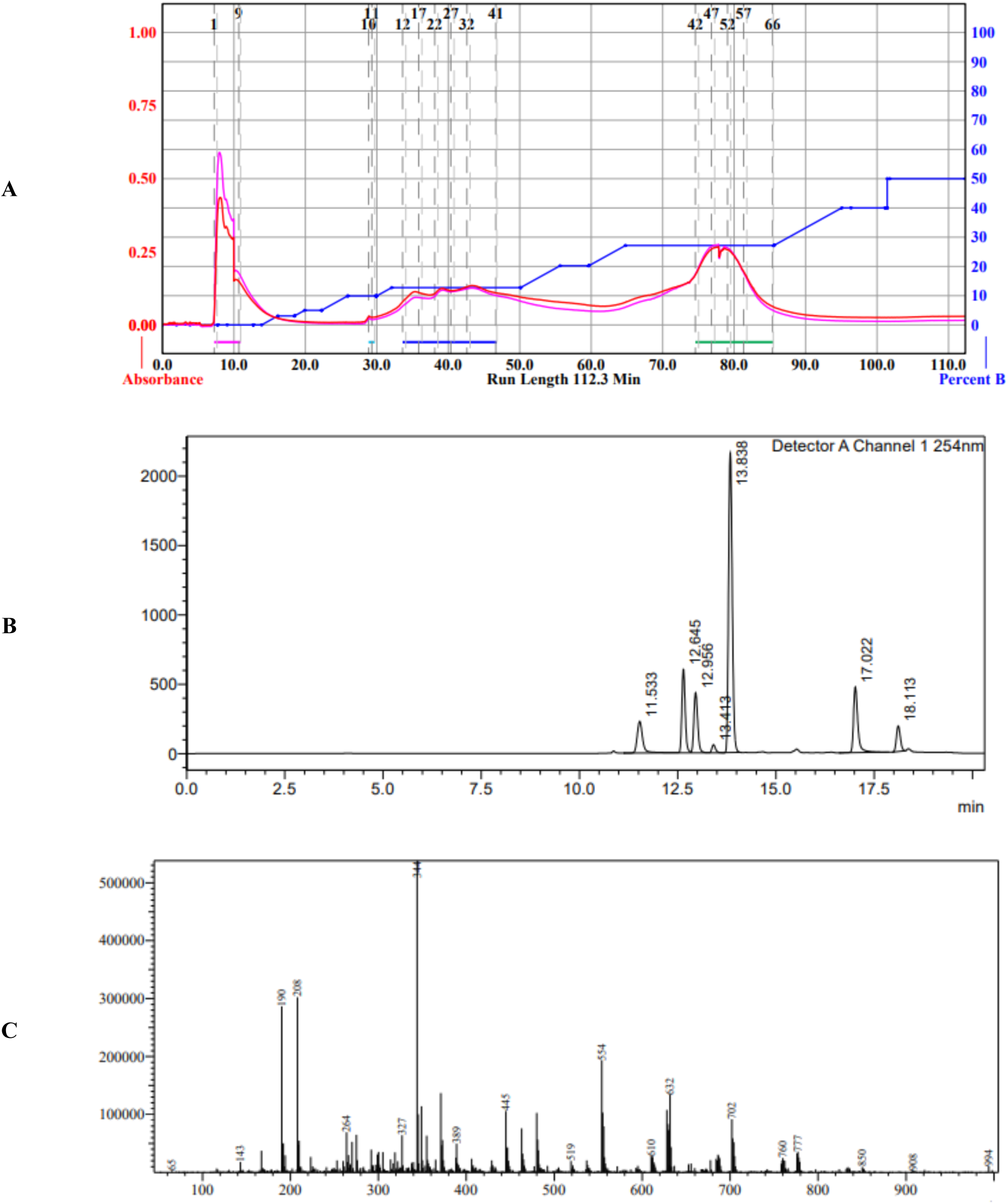
Chromatograms and spectra of CI plant extracts, **A:** Flash chromatogram of the fractions in CI methanol extract, **B:** HPLC chromatogram of the fractions in CI methanol extract, **C:** LC-MS spectra fo

**Figure 9:**
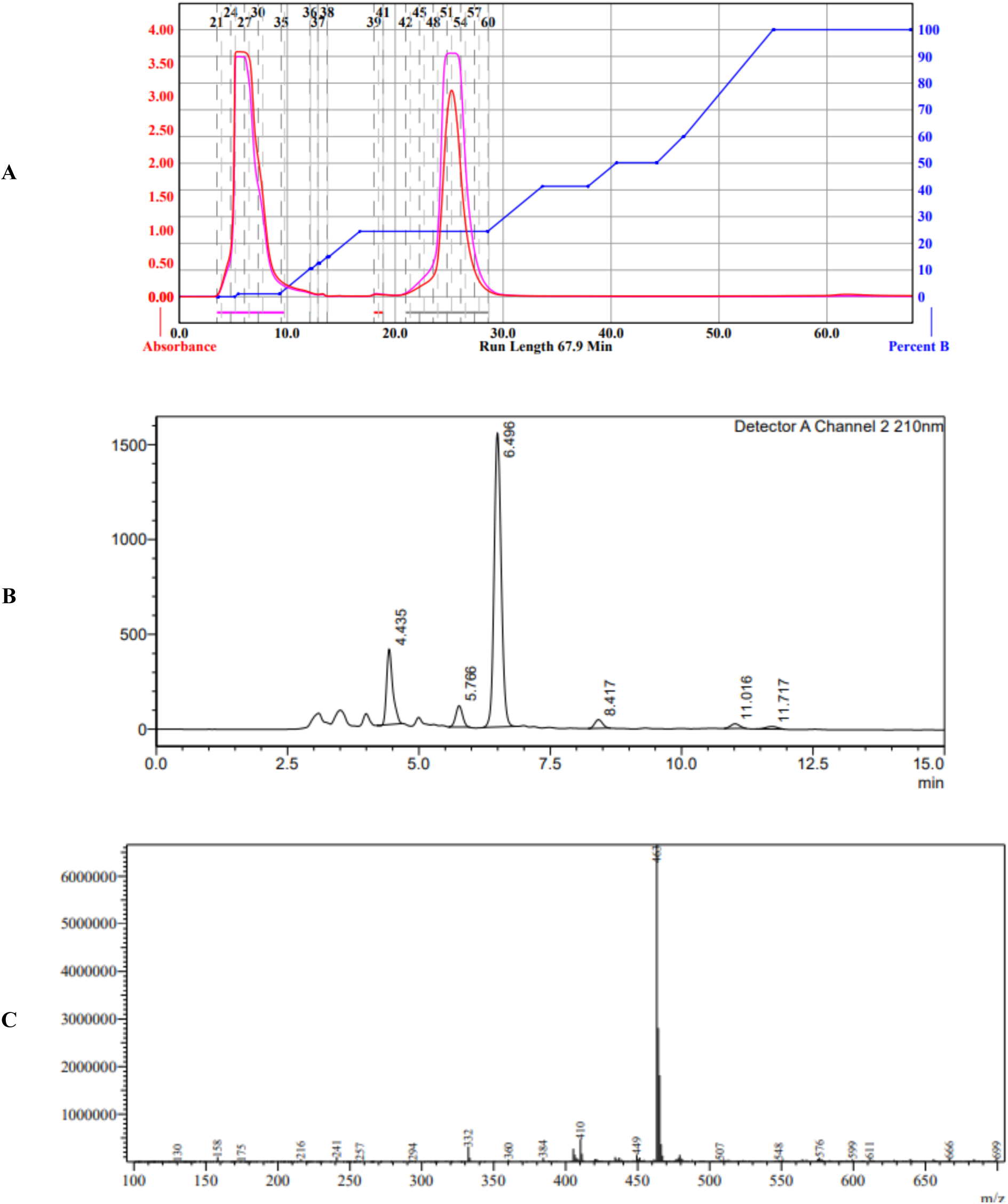
Chromatograms and spectra of AP plant extracts, **A:** Flash chromatogram of the fractions in AP methanol extract, **B:** HPLC chromatogram of the fractions in AP methanol extract, **C:** LC-MS spectra

**Figure 9:**
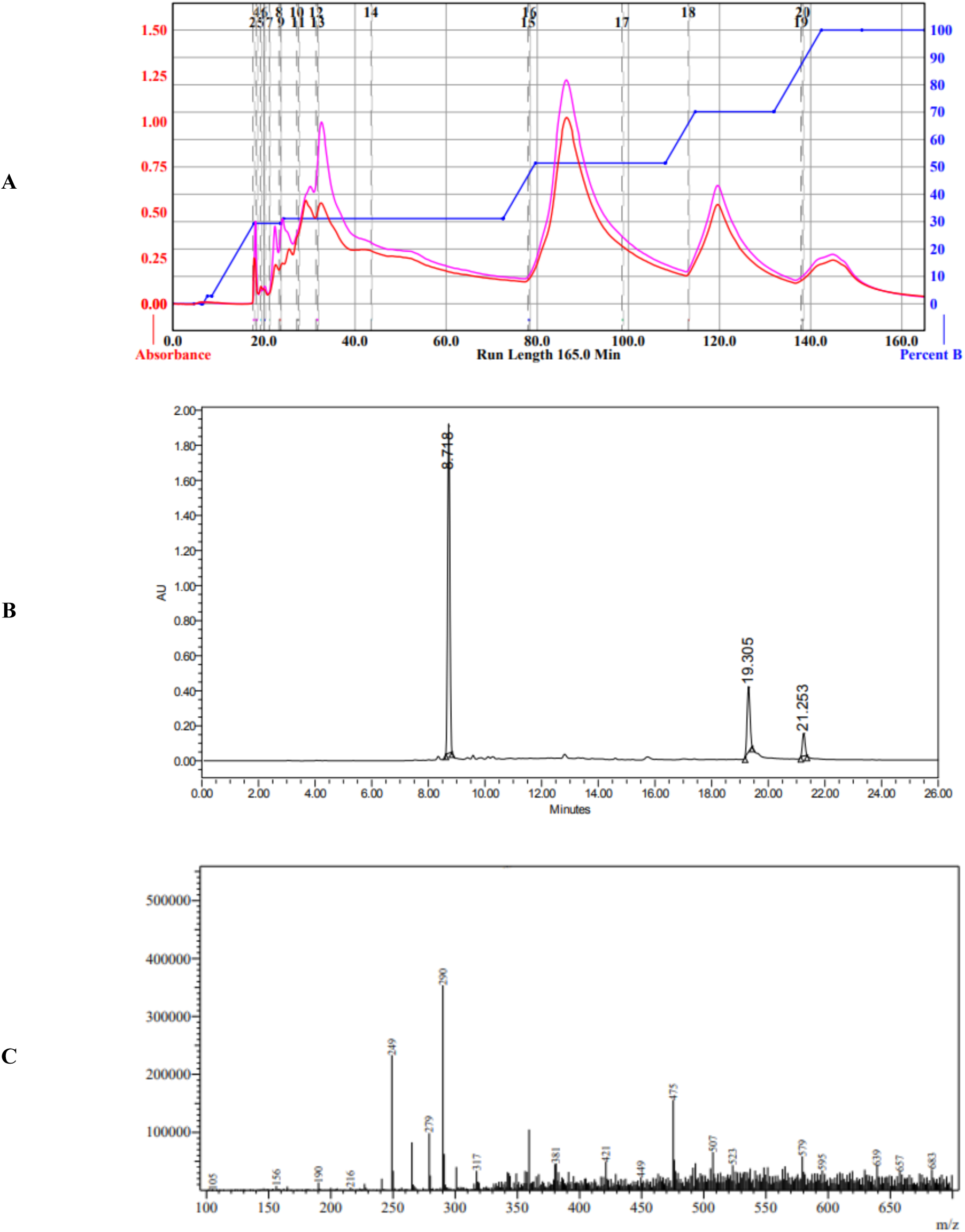
Chromatograms and spectra of AP plant extracts, **A:** Flash chromatogram of the fractions in AP methanol extract, **B:** HPLC chromatogram of the fractions in AP methanol extract, **C:** LC-MS spectra

The flash chromatographic fractionation of crude methanol extracts of AP has led to two different polarity fractions (**Figure 9A**) at different RT (**Table 8**). The HPLC chromatogram for the second fraction obtained from flash chromatography at Rt2 of 25min has shown a high resolution peak at the retention time of 6.5min (**Figure 9B**) based on UV wavelength at 210 nm. The HPLC fraction obtained at the retention time of 6.5min has shown a peak corresponding 463 m/z in MS spectra (**Figure 9C**) has confirmed the presence of Andrographidine C in AP methanol extract. Hence, it is inferred that the Andrographidine C in the extract can interact with RdRp of SARS-CoV-2 and offer the Anti-SARS-CoV-2 activity.

The flash chromatographic fractionation of crude methanol extracts of CA has led to four different polarity fractions (**Figure 10A**) at different RT (**Table 8**). The HPLC chromatogram for the second fraction obtained from flash chromatography at Rt2 of 30min has shown a high resolution peak at the retention time of 8.71min (**Figure 10B**) based on UV wavelength at 370nm. The HPLC fraction obtained at the retention time of 8.71min has shown a peak corresponding 290 m/z in MS spectra (**Figure 10C**) has confirmed the presence of Kaempferol C in CA methanol extract. Hence, it is inferred that, the Kaempferol in the extract can interact with S-Protein of SARS-CoV-2 and offer the Anti-SARS-CoV-2 activity.

**Figure 10:**
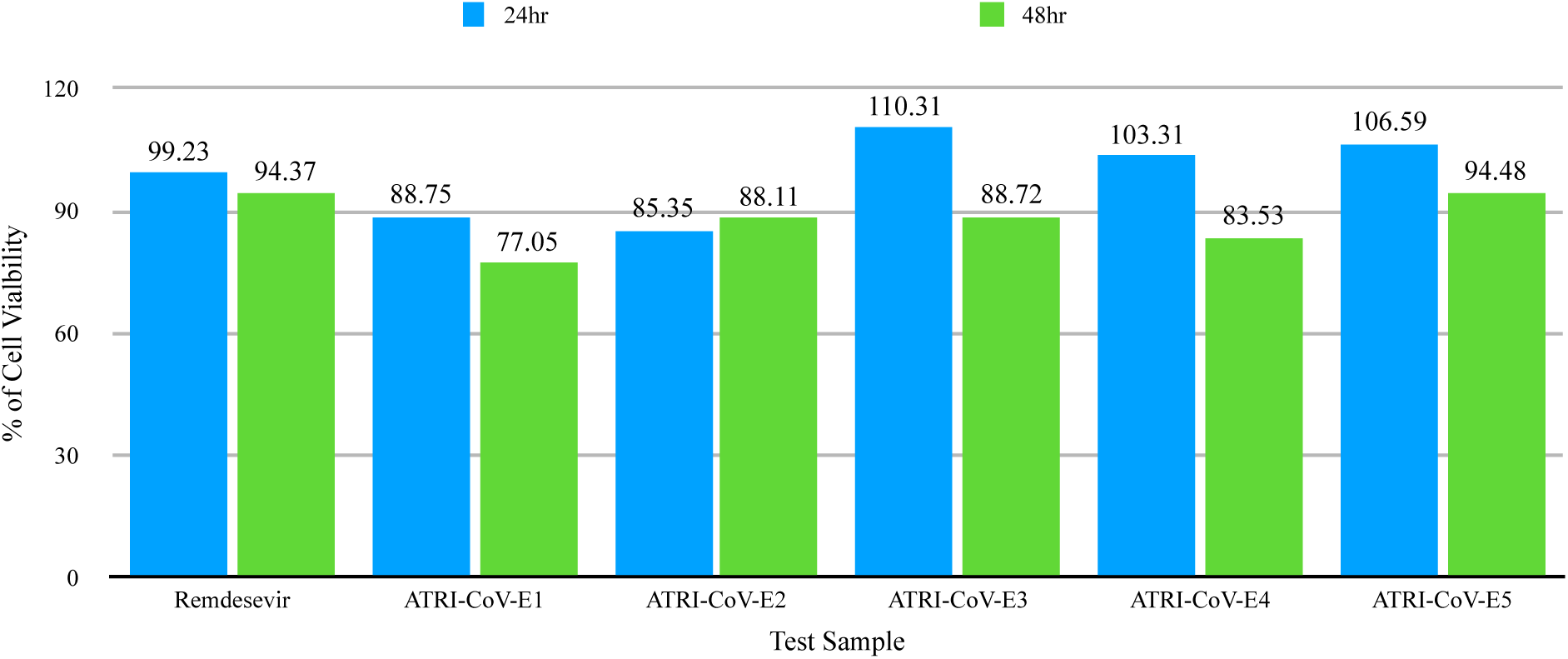
The viability of VeroE6 cells treated with plant extracts and Remdesevir for 24 and 48hr.

### 3.3. *In vitro* cytotoxicity

The extracts ATRI-CoV-E3, ATRI-CoV-E4 and ATRI-CoV-E5 showed promotion of VeroE6 cell growth with 110.31%, 103.31% and 106.59% cell viability at 24hr post treatment respectively. ATRI-CoV-E3, ATRI-CoV-E4 and ATRI-CoV-E5 showed 88.72%, 83.53% and 94.48% of cell viability at 48hr post treatment. The extracts ATRI-CoV-E1 and ATRI-CoV-E2 showed 88.75% and 77.05%, 85.35% and 88.11% of cell viability at 24hr and 48hr post treatment respectively. Remdesivir at 10µM concentration showed 99.23% and 94.37 % cell viability at 24 and 48 hr respectively as shown in **Figure 10**. TS showed greater cell viability after 48 hr of treatment as compared to Remdesivir.

### 3.4. *In vitro* Anti-SARS-CoV-2 Activity

For E gene, at 24 hr, Remdesevir, ATRI-CoV-E1, ATRI-CoV-E2, ATRI-CoV-E3, ATRI-CoV-E4 and ATRI-CoV-E5 showed 82.28%, 47.86%, 45.93%, -31.50%, 73.54%, and 74.75% inhibition respectively. For the N gene, the inhibition was 80.35%, 49.50%, 45.47%, -19.20%, 71.95%, 74.50% respectively (**Figure 11**).

**Figure 11:**
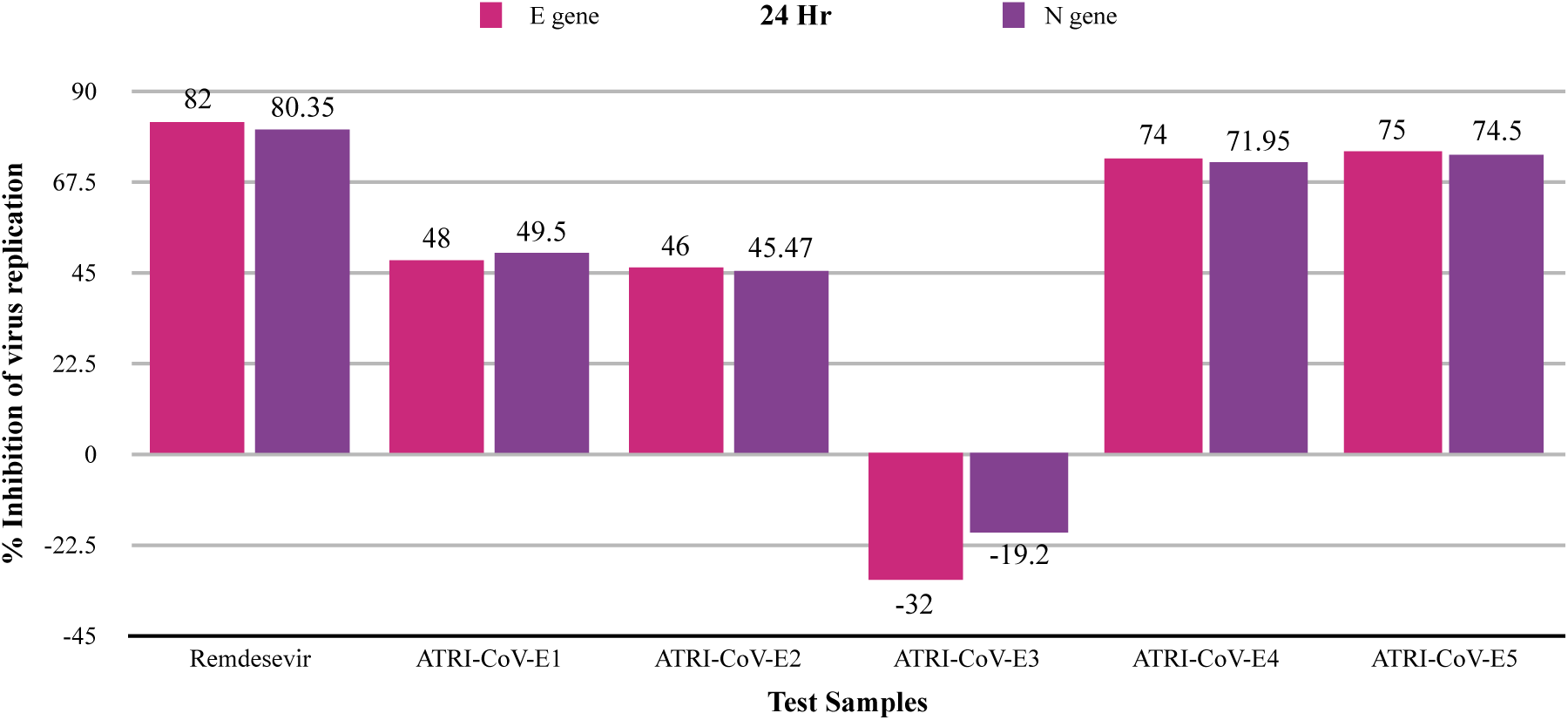
Percentage inhibition of SARS-CoV-2 replication by Remdesevir and TSs after 24hr of treatment considering the replication of E and N genes.

For E gene, at 48 hr, Remdesevir, ATRI-CoV-E1, ATRI-CoV-E2, ATRI-CoV-E3, ATRI-CoV-E4 and ATRI-CoV-E5 showed 99.80%, 91.30%, 95.60%, 71.72%, 99.90% and 92.07% inhibition. For the N gene, the inhibition was 99.89%, 81.14%, 91.50%, 55.90%, 99.80% and 84.64% respectively (**Figure 12**).

**Figure 12:**
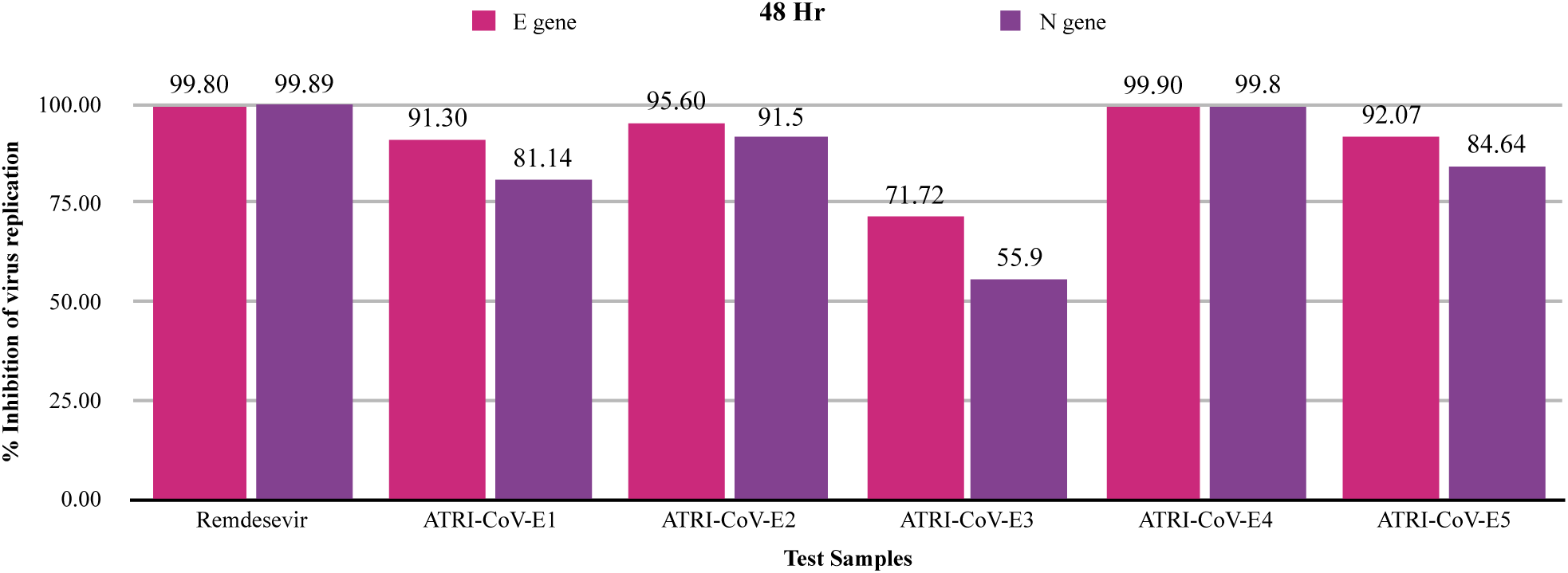
Percentage inhibition of SARS-CoV-2 replication by Remdesevir and TSs after 48hr of treatment considering the replication of E and N genes.

Considering the outcomes of *in vitro* Anti-SARS-CoV-2 activity, the plant extracts ATRI-CoV-E2, ATRI-CoV-E4 and ATRI-CoV-E5 were subjected for seven-point IC_50_ estimation to identify the minimum dose required to inhibit the replication of SARS-CoV-2.

### 3.5. *In vitro* IC_50_ determination

The dose-response curves were obtained using seven different concentrations of Remdesivir, ATRI-CoV-E2, ATRI-CoV-E4, and ATRI-CoV-E5. For E gene and N gene, Remdesivir showed IC_50_ of 0.15 µM and 0.11 µM respectively (**Figure 13A and 13B)**, For E gene and N gene, ATRI-CoV-E4 showed IC_50_ of 1.18 µg and 1.16 µg respectively (**Figure 14A and 14B)**. Remdesivir and ATRI-CoV-E4 showed significant delta CT for both E and N genes as shown in **Table 9**. The Delta CT of ATRI-CoV-E2 and ATRI-CoV-E5 were moderate. In IC_50_ estimation, ATRI-CoV-E2 and ATRI-CoV-E5 did not show consistent CT values, which is common to see in replication of experiments (Vaux et al. 2012). However, the anti-SARS-CoV2 activity of ATRI-CoV-E2 and ATRI-CoV-E5 were significant. The anti-viral activity of *W. somnifera* (ATRI-CoV-E2) and *C. asiatica* (ATRI-CoV-E5) has also been reported by earlier researchers on other viruses besides SARS-CoV-2 (Kashyap VK et al. 2020 and Sun et al. 2020).

**Figure 13:**
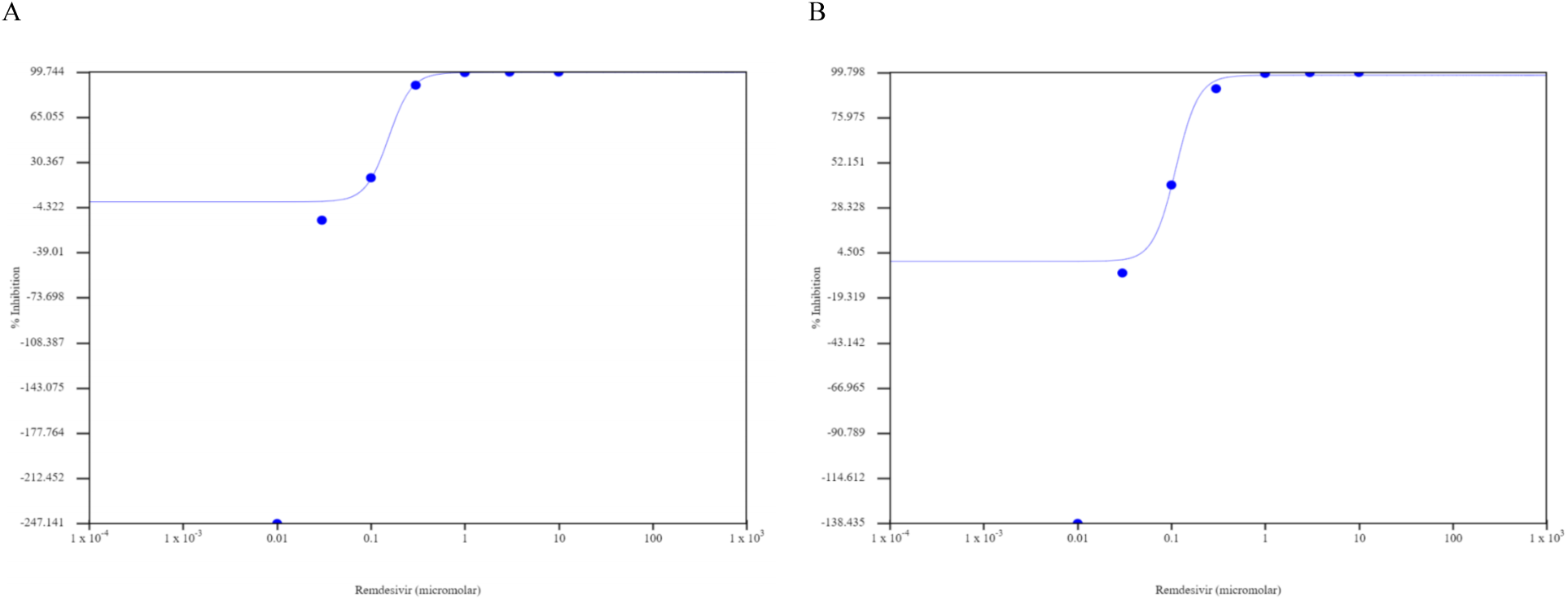
(A) Dose dependent response of Remdesivir for SARS-CoV-2 E gene (B) Dose dependent response of Remdesivir for SARS-CoV-2 N gene.

**Figure 14:**
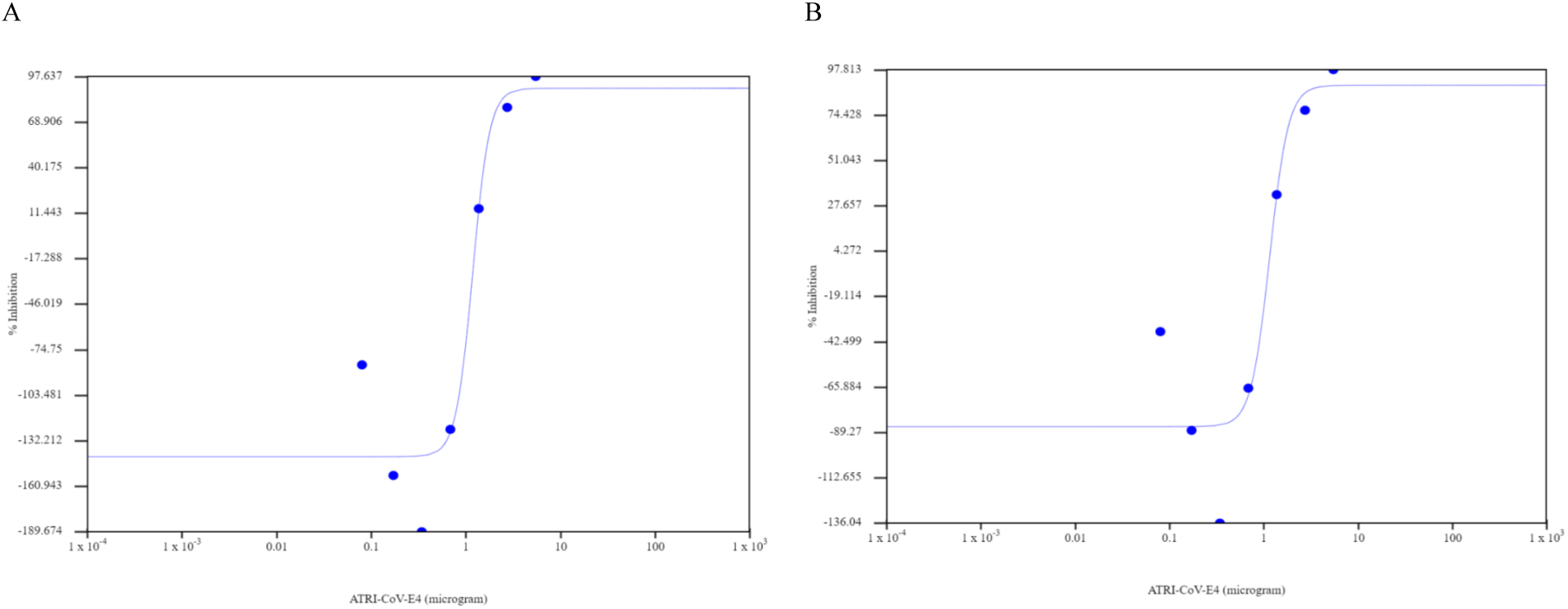
(A) Dose dependent response of ATRI-CoV-E4 for SARS-CoV-2 E gene (B) Dose dependent response of ATRI-CoV-E4 for SARS-CoV-2 N gene.

**Table 9:** Summary of data relating to SARS-CoV-2 real-time PCR Ct value in the presence and absence of control and TSs

| E GENE- 48 hours |  | Ct Mean |  |  |  |  |  |  |  |
| --- | --- | --- | --- | --- | --- | --- | --- | --- | --- |
| S. No. | Sample Name | #1 | #2 | #3 | Average CT | Delta CT | Fold change | wrt control | % Inhibition |
| 1 | DMSO | 14.780 | 13.974 | 14.169 | 14.308 | -0.000 | 1.000 | 100.0 | -0.0 |
| 2 | REMDE-SIVIR | 25.392 | 25.316 | 25.123 | 25.277 | 10.969 | 2004.441 | 0.050 | 99.950 |
| 3 | ATRI-CoV-E2 | 19.201 | 19.151 | 18.056 | 18.803 | 4.495 | 22.545 | 4.44 | 95.6 |
| 4 | ATRI-CoV-E4 | 24.554 | 23.495 | 23.325 | 23.791 | 9.483 | 715.69 | 0.14 | 99.9 |
| 5 | ATRI-CoV-E5 | 18.956 | 17.025 | 17.913 | 17.964 | 3.656 | 12.61 | 7.93 | 92.070 |
| N GENE- 48 hours |  | Ct Mean |  |  |  |  |  |  |  |
| S. No. | Sample Name | #1 | #2 | #3 | Average CT | Delta CT | Fold change | wrt control | % Inhibition |
| 1 | DMSO | 15.165 | 14.322 | 14.659 | 14.715 | 0.000 | 1.000 | 100.0 | 0.0 |
| 2 | REMDE-SIVIR | 24.895 | 24.801 | 24.419 | 24.705 | 9.990 | 1016.882 | 0.098 | 99.902 |
| 3 | ATRI-CoV-E2 | 18.673 | 18.251 | 17.897 | 18.274 | 3.559 | 11.785 | 8.5 | 91.5 |
| 4 | ATRI-CoV-E4 | 24.535 | 22.997 | 22.817 | 23.450 | 8.735 | 425.992 | 0.2 | 99.8 |
| 5 | ATRI-CoV-E5 | 18.290 | 16.666 | 17.300 | 17.418 | 2.703 | 6.514 | 15.35 | 84.648 |

There is a consensus among virologists that combining two or more therapeutic agents with antiviral activity could result in improved antiviral response (Richman et al. 2016; Balkrishna et al. 2021). We therefore decided to evaluate the *in vivo* toxicity of combined extracts of ATRI-CoV-E2, ATRI-CoV-E4 and ATRI-CoV-E5 (ATRICOV 452).

### 3.4. In vivo toxicity

#### 3.4.1. Acute toxicity

No abnormalities were observed in clinical signs as shown in **Table 10**. Body weights of each animal were recorded prior to the administration of ATRICOV 452 (Day 0) and at the end of the experiment (Day 15). The highest dose of ATRICOV 452 2000 mg/kg body weight did not induce acute toxicity in rats during the study and the ATRICOV 452 fed animals showed healthy growth during the observation period as shown in **Table 11**.

**Table 10:**
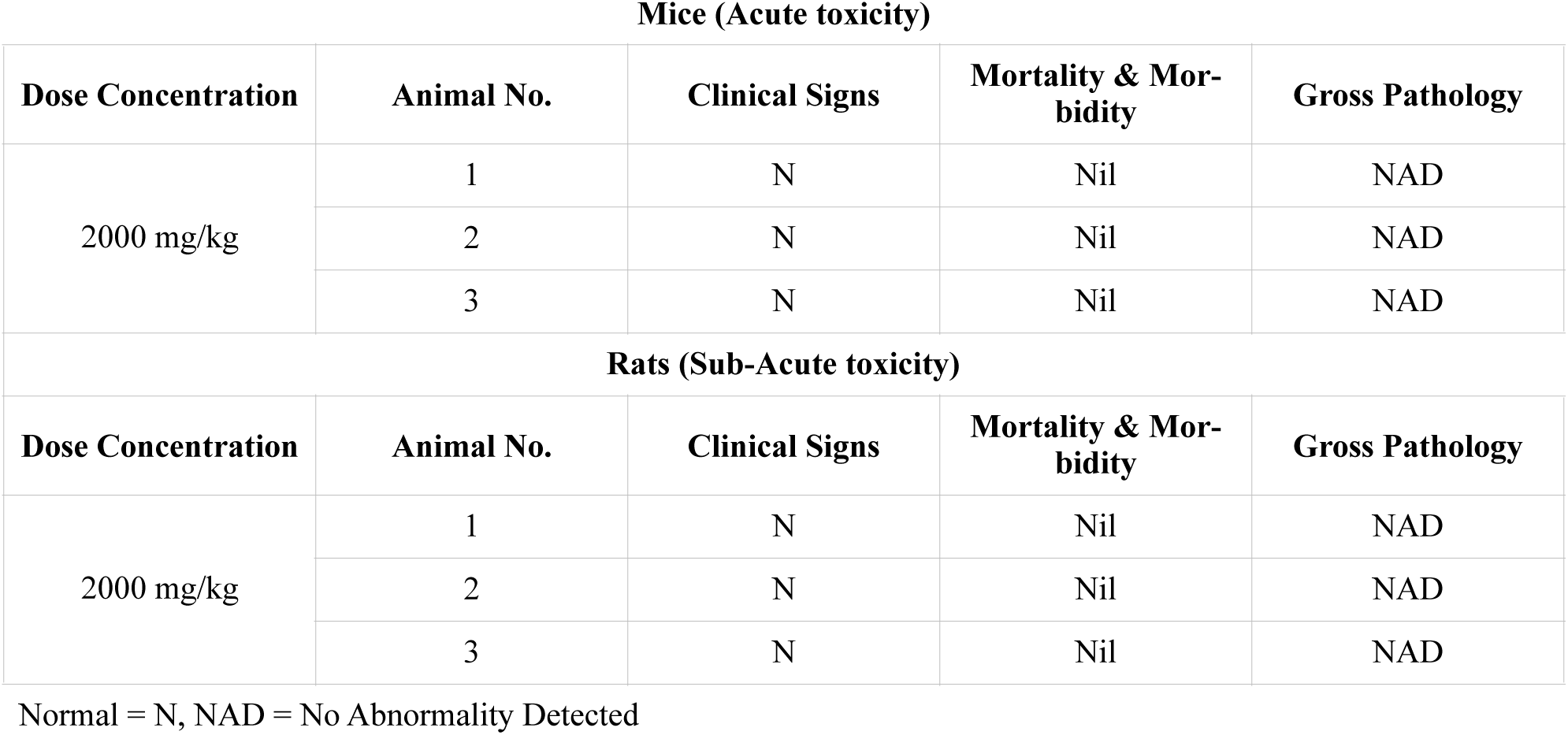
Results of Clinical signs, Mortality & Morbidity, and Gross pathology

**Table 11:**
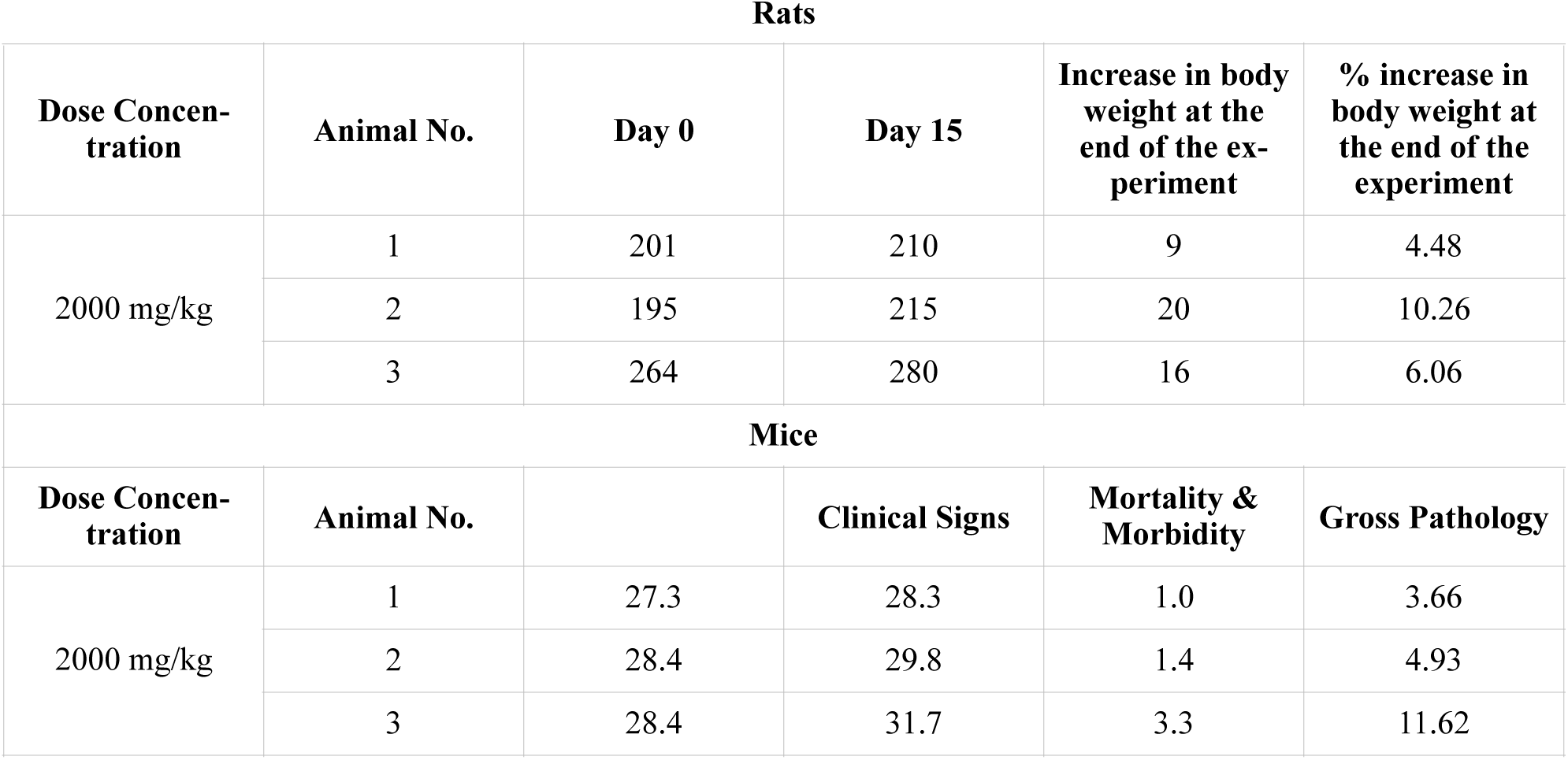
Individual animal body weights (g) in acute toxicity

#### 3.4.2. Sub-acute toxicity

During the study period, clinical signs of all treated animals were found normal as tabulated in **Table 12**. The food intake and body weights increased as shown in **Table 13**.

**Table 12:** Summary of Clinical Signs and Mortality of animals

| **Group/Dose (mg/kg)** | **Male** |  | **Female** |  |
| --- | --- | --- | --- | --- |
|  | **Clinical Sign** | **Mortality** | **Clinical Sign** | **Mortality** |
| GI (0.0) | N | 0 | N | 0 |
| GII (200.0) | N | 0 | N | 0 |
| GIII (400.0) | N | 0 | N | 0 |
| GIV (800.0) | N | 0 | N | 0 |
| GV (0.0) | N | 0 | N | 0 |
| GVI (800.0) | N | 0 | N | 0 |
| Abbreviations': N - Normal, 0 - No Mortality, E-Emaciation |  |  |  |  |

**Table 13:** Summary of Group Mean Body Weight of animals in sub-acute toxicity study

|  | Male animals |  |  |  |  |  |  |
| --- | --- | --- | --- | --- | --- | --- | --- |
| Study Day | Group | Control | Low | Medium | High | Control Reversal | High Reversal |
| 1 | Mean | 178.33 | 151.67 | 150.00 | 190.83 | 134.17 | 177.50 |
|  | ± SD | 11.25 | 17.22 | 16.73 | 11.14 | 13.20 | 24.85 |
|  | n | 6 | 6 | 6 | 6 | 6 | 6 |
| 8 | Mean | 190.00 | 168.33 | 188.33 | 210.00 | 150.83 | 180.83 |
|  | ± SD | 12.25 | 21.37 | 22.73 | 14.14 | 13.93 | 25.18 |
|  | n | 6 | 6 | 6 | 6 | 6 | 6 |
| 15 | Mean | 188.33 | 206.67 | 208.33 | 233.33 | 167.50 | 197.50 |
|  | ± SD | 9.83 | 21.37 | 24.01 | 13.29 | 13.69 | 21.62 |
|  | n | 6 | 6 | 6 | 6 | 6 | 6 |
| 22 | Mean | 210.83 | 217.50 | 224.17 | 234.17 | 185.83 | 220.00 |
|  | ± SD | 9.17 | 22.30 | 23.54 | 10.68 | 17.72 | 23.45 |
|  | N | 6 | 6 | 6 | 6 | 6 | 6 |
| 29 | Mean | 231.67 | 238.33 | 249.17 | 235.00 | 191.67 | 215.83 |
|  | ± SD | 16.02 | 22.73 | 26.91 | 14.49 | 14.72 | 22.23 |
|  | N | 6 | 6 | 6 | 6 | 6 | 6 |
| 36 | Mean | NA |  |  |  | 196.67 | 214.17 |
|  | ± SD |  |  |  |  | 17.51 | 28.18 |
|  | N |  |  |  |  | 6 | 6 |
| 43<br>(Pre-fasting) | Mean | NA |  |  |  | 215.00 | 224.17 |
|  | ± SD |  |  |  |  | 16.12 | 36.25 |
|  | N |  |  |  |  | 6 | 6 |
|  | Female animals |  |  |  |  |  |  |
| Study Day | Group | Control | Low | Medium | High | Control Reversal | High Reversal |
| 1 | Mean | 145.00 | 138.33 | 143.33 | 161.67 | 152.50 | 160.00 |
|  | ± SD | 8.37 | 18.62 | 15.38 | 9.83 | 13.32 | 15.17 |
|  | n | 6 | 6 | 6 | 6 | 6 | 6 |
| 8 | Mean | 152.50 | 144.17 | 150.00 | 166.67 | 162.50 | 173.33 |
|  | ± SD | 14.05 | 23.33 | 16.43 | 6.06 | 12.14 | 16.63 |
|  | n | 6 | 6 | 6 | 6 | 6 | 6 |
| 15 | Mean | 165.83 | 155.00 | 162.50 | 173.33 | 166.67 | 181.67 |
|  | ± SD | 14.63 | 24.08 | 19.43 | 5.16 | 10.33 | 19.41 |
|  | n | 6 | 6 | 6 | 6 | 6 | 6 |
| 22 | Mean | 172.50 | 166.67 | 171.67 | 178.33 | 178.33 | 186.67 |
|  | ± SD | 16.36 | 25.82 | 19.41 | 14.72 | 6.83 | 16.33 |
|  | **n** | 6 | 6 | 6 | 6 | 6 | 6 |
| **29** | **Mean** | **177.50** | **175.83** | **182.50** | **180.00** | **175.00** | **187.50** |
|  | **± SD** | 19.17 | 22.23 | 20.19 | 14.83 | 15.81 | 13.32 |
|  | **n** | 6 | 6 | 6 | 6 | 6 | 6 |
| **36** | **Mean** | NA |  |  |  | **179.17** | **183.33** |
|  | **± SD** |  |  |  |  | 13.93 | 17.22 |
|  | **n** |  |  |  |  | 6 | 6 |
| **43**<br>**(Pre-fasting)** | **Mean** | NA |  |  |  | **180.83** | **187.50** |
|  | **± SD** |  |  |  |  | 14.97 | 19.17 |
|  | **n** |  |  |  |  | 6 | 6 |

##### 3.4.2.a. Haematological analysis

The administrations of ATRICOV 452 in high dose showed increase in RBC and reticulocyte count in both male (8.707 × 10^6^/mm3 Cells ) and female (7.278 × 10^6^/mm3) compared with the control (4.75 × 10^6^/mm3) however it was not statistically significant (**Figure 15 and Figure 16**).

**Figure 15:**
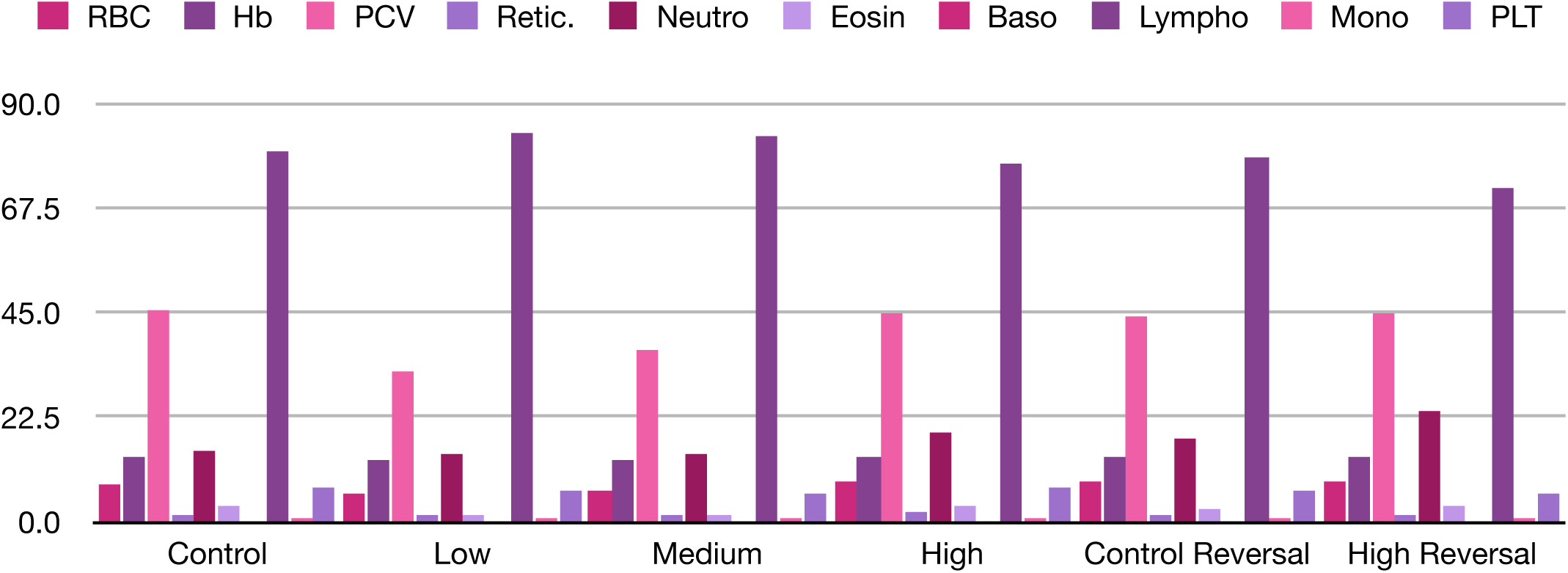
Summary of group mean hematological estimations of male rats treated with different doses.

**Figure 16:**
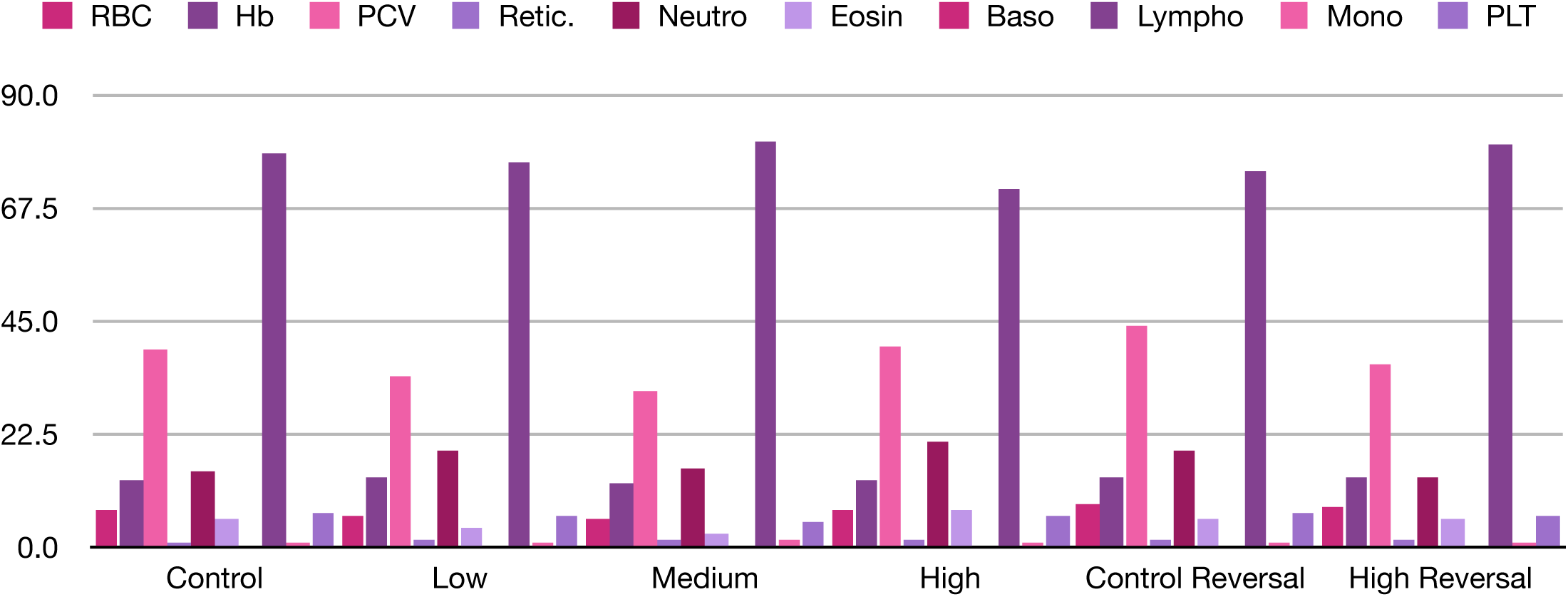
Summary of group mean hematological estimations of female rats treated with different doses.

PCV, platelet count and eosinophil count in male (Low dose: 1.16 × 10^3^/mL and Medium dose: 1.16 × 10^3^/mL) and female (Low dose: 3.50 × 10^3^/mL and Medium dose: 2.33 × 10^3^/mL) animals showed statistically significant (P<0.05) decrease compared to control group (**Figure 15 and Figure 16)**. Rest of the hematological parameters (Haemoglobin, Neutrophils, Basophils, Lymphocytes and Monocytes101010) showed no change (**Figure 15 and Figure 16**).

##### 3.4.2.b. Biochemical analysis

Biochemical analysis was conducted on day 29 and 43; the following observations were made by comparing with control groups.

• *Glucose* The male animals receiving high dose treatment showed statistically significant (P<0.05) decrease in glucose. The low and medium dose groups showed no statistically significant change (**Figure 17)**.
• *Aspartate Aminotransferase (AST)* Male animals receiving low, medium and high dose treatment showed significant (P<0.0001, P<0.01, P<0.001) increase in AST (**Figure 17**). However, such changes were not evident in female animals (**Figure 18**).
• *Alkaline Phosphatase (ALP)* Male animals receiving medium and high dose showed significant (P<0.001) decrease in ALP (**Figure 17)**.
• *Total protein* Male and animals receiving low, medium and high dose treatment showed statistically significant (P<0.05) increase in total protein value (**Figure 17 and Figure 18)**.
• *Total cholesterol* Male animals receiving low and medium dose treatment showed significant (P<0.05) increase in total cholesterol (**Figure 17)**.
• *Triglyceride* Female animals receiving low and high dose treatment showed significant (P<0.05) increase in triglyceride (**Figure 18**).
• *Blood Urea Nitrogen* Female animals receiving medium dose treatment showed significant ( P<0.01) decrease in BUN (**Figure 18**).
• *Creatinine* Male and female animals receiving low and medium dose treatment showed significant (P<0.0001) decrease in creatinine (**Figure 17 and Figure 18)**.
• *Total bilirubin* Female animals receiving high dose treatment show significant (P<0.05) decrease in total bilirubin (**Figure 18**).
• *Phosphorous* Male animals receiving low dose treatment showed significant (P<0.01) decreases in phosphorus (**Figure 17**).

**Figure 17:**
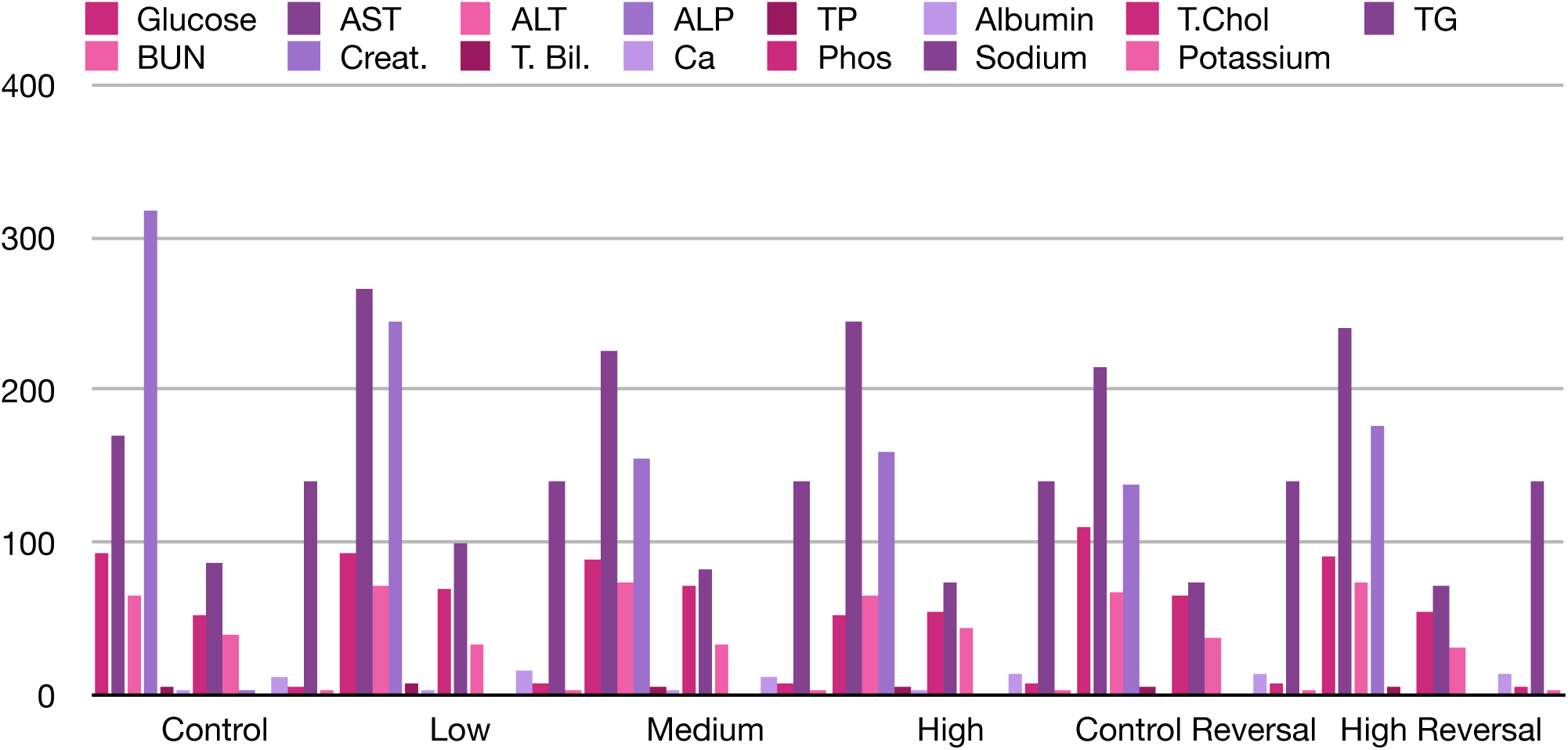
Summary of group mean clinical chemistry estimations of male rats treated with different doses.

**Figure 18:**
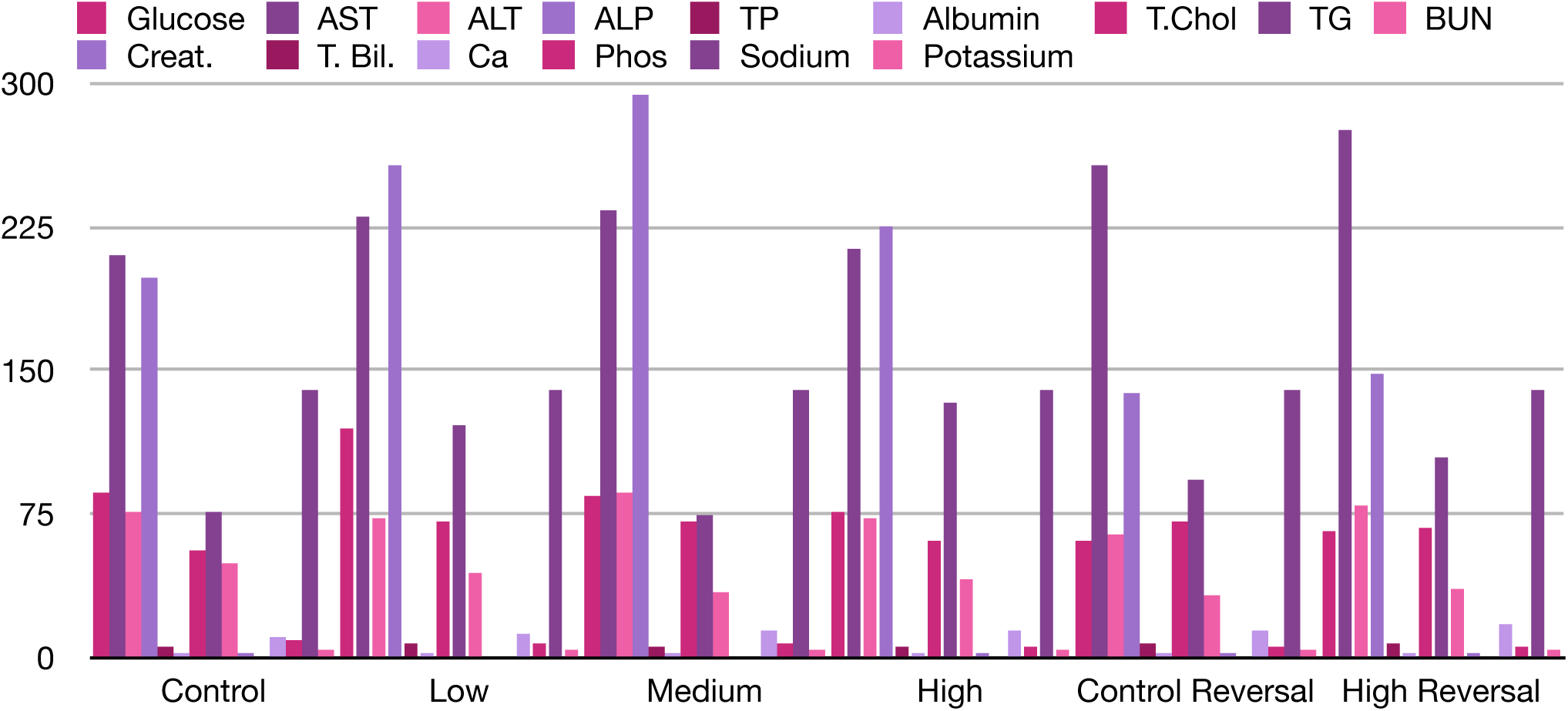
Summary of group mean clinical chemistry estimation of female rats treated with different doses.

##### 3.4.2.c. Bone marrow analysis

Bone marrow analysis was conducted on day 29 and day 43. At low dose, treatment groups showed mild reactive plasmacytosis. At medium dose, treatment groups showed mild reactive myeloid hyperplasia. At high dose, treatment groups showed mild to moderate myeloid hyperplasia and mild reactive plasmacytosis. The control groups showed no bone marrow change. (**Table 14)**.

**Table 14:** Summary of bone marrow study performed in male and female animals treated with ATRICOV 452 in sub-acute toxicity study

| Group (Dose in mg/kg) | Observation |
| --- | --- |
| **GI (0.0)** | All the six bone marrow smears from control group 6CM1 to 6CM6 show features suggestive of normal haematopoiesis.<br>All the six bone marrow smears from the second control group 6CF1 to 6CF6 show features suggestive of normal haematopoiesis. |
| **GII (200.0)** | All the six bone marrow smears in the Low Dose group from 6LM1 to 6LM6 and 6LF1 to 6LF6 show features of MILD REACTIVE PLASMACYTOSIS.<br>No pathological changes seen. |
| **GIII (400.0)** | All the six bone marrow smears in the medium dose group form 6MM1 to 6MM6 and 6MF1 to 6MF6 show features indicative of MILD REACTIVE MYELOID HYPERPLASIA.<br>No specific pathological changes seen. |
| **GIV (800.0)** | All the six bone marrow smears in the high dose group from 6HM1 to 6HM6 and 6HF1 to 6HF6 show features of MILD TO MODERATE MYELOID HYPERPLASIA AND MILD REACTIVE PLASMACYTOSIS.<br>No specific pathological changes seen. |
| **GV (0.0)** | All the six bone marrow smears in the reversal control group 6CRM1 to 6CRM6 and 6CRF1 to 6CRF6 show features of NORMAL HAEMATOPOIESIS.<br>No pathological changes seen. |
| **GVI (800.0)** | All the six bone marrow smears in the reversal high dose control group 6HRM1 to 6HRM6 and 6HRF1 to 6HRF6 show features indicative of NORMAL HAEMATOPOIESIS / MILD REACTIVE PLASMACYTOSIS.<br>No other haematopoietic abnormalities seen in any of these smears in the reversal groups. |

Abbreviations: 6CM1- 6 control male 1, where 6 is total number of animals in group and 1 is an animal in group. CF- control female, LM- Low dose male, LF- Low dose female, MM- Medium dose male, LF- Medium dose female, HM- High dose male, HF- High dose female, CRM - Control reversible male, CRF - Control reversible female, HRM - High dose reversible male, HRF - High dose reversible female

##### 3.4.2.d. Urine analysis

Urine analysis showed no changes on day 29 and on day 43.

##### 3.4.2.e. Organ weight

The organ weight comparison was done with control groups. Liver weight of male animals receiving high dose treatment showed significant (P<0.05) decrease. Spleen of male animals receiving low dose and female animals receiving medium dose showed significant (P<0.05) increase in weight. Lungs of female animals receiving medium dose showed significant (P<0.05) decrease (**Table 15**).

**Table 15:**
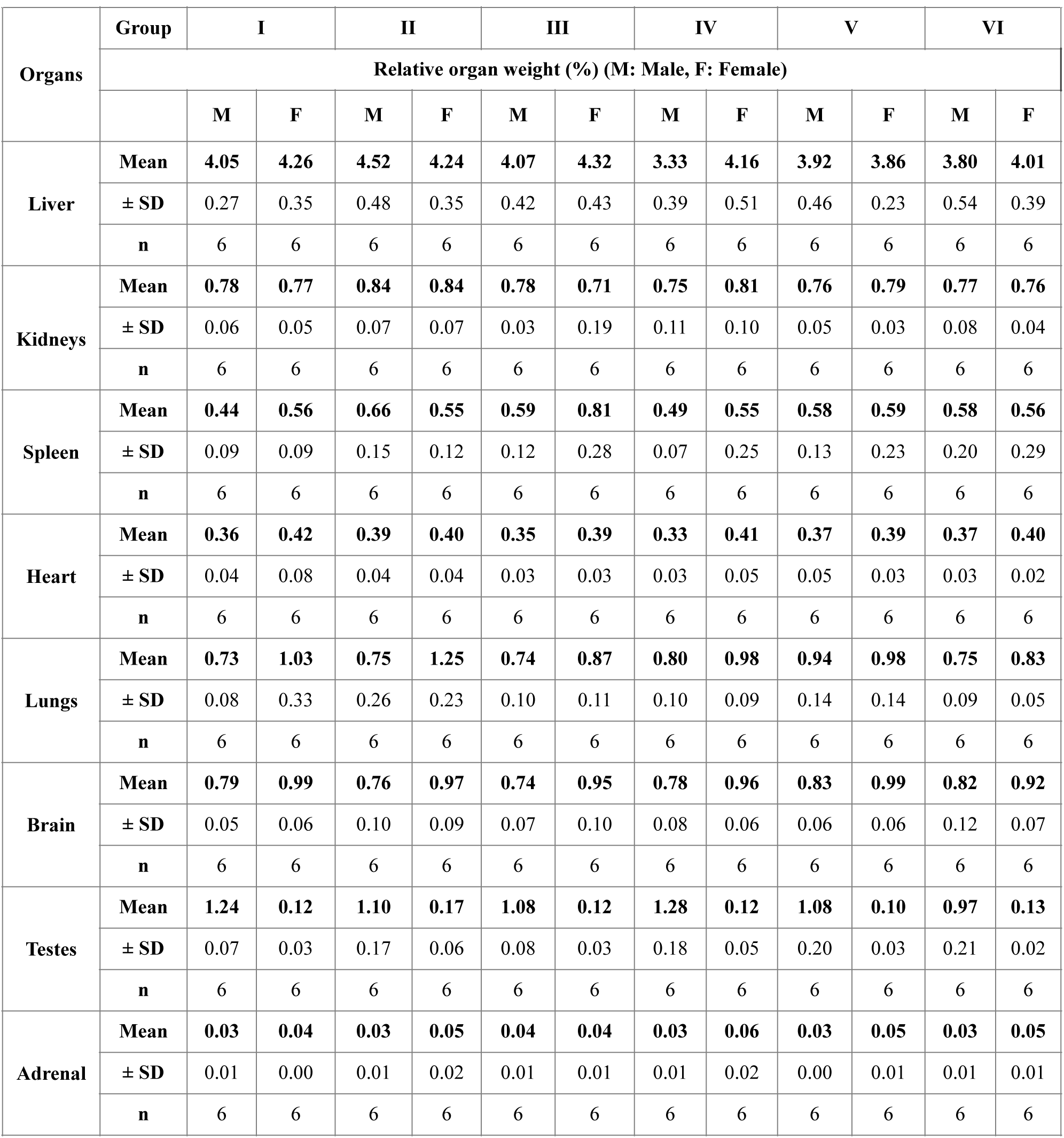
Summary of group mean relative organ weight in male and female animals treated with ATRICOV 452 in sub-acute toxicity study

For the rest of the organs (Kidneys, Heart, Brain, Testes and Adrenal) no changes in the weight was observed.

##### 3.4.2.f. Histopathology

No histopathological changes were observed in the liver, kidney, and jejunum of animals treated with low, medium and high dose (**Figures 19**, **Figures 20 and Figures 21)**.

**Figure 19:**
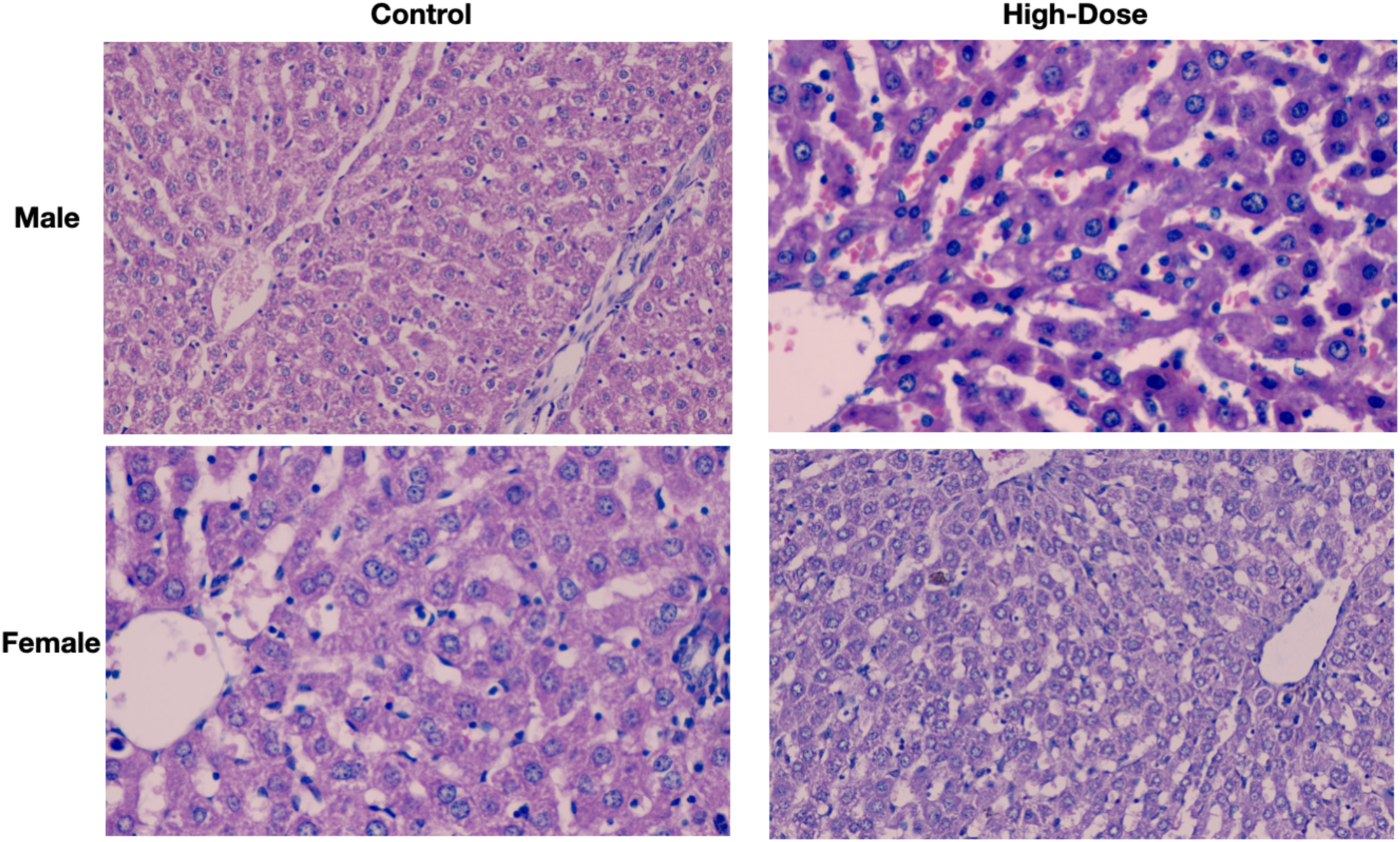
Histopathological analyses of the liver in high-dose male and female rats with/without ATRICOV 452 treatment.

**Figure 20:**
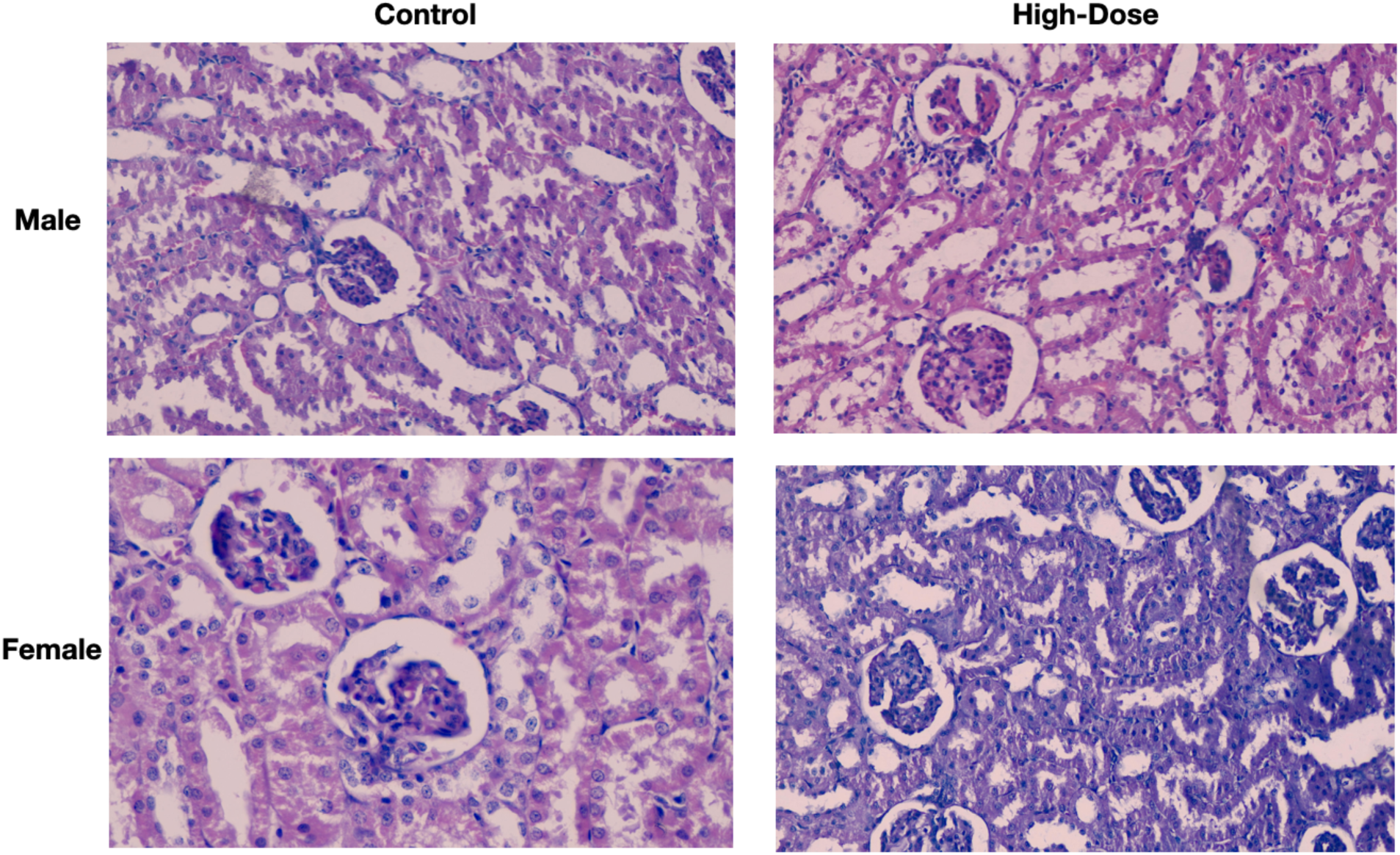
Histopathological analyses of the kidney in high-dose male and female rats with/without ATRICOV 452 treatment.

**Figure 21:**
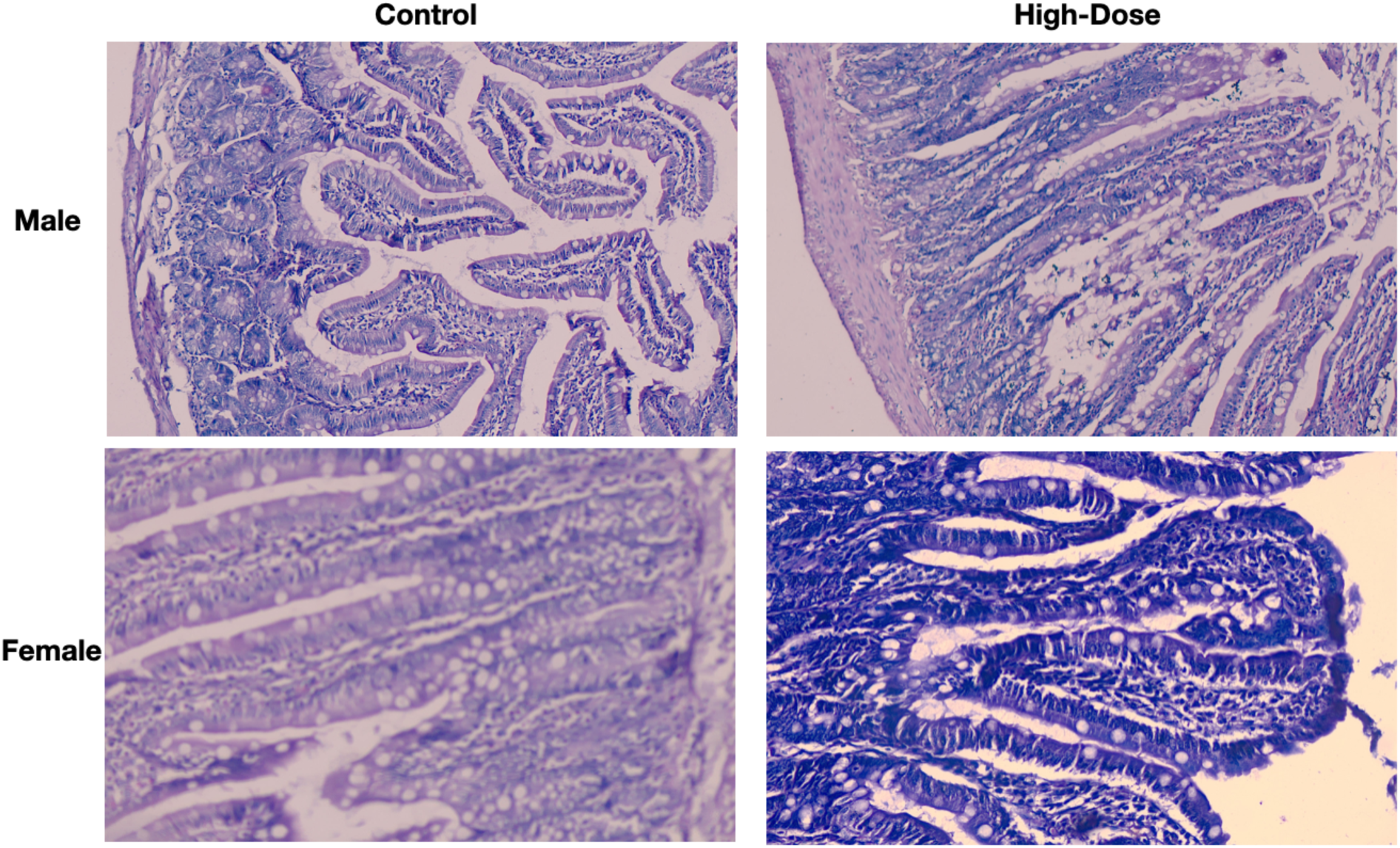
Histopathological analyses of the jejunum in high-dose male and female rats with/without ATRICOV 452 treatment.

## 4. Conclusions

We have demonstrated that the *in vitro* Anti-SARS-CoV-2 activity of plant extracts ATRI-CoV-E4, ATRI-CoV-E5, and ATRI-CoV-E2, showed no cytotoxicity in VeroE6 cells. The seven-point IC_50_ study of extracts showed consistent Anti-SARS-CoV-2 activity of ATRI-CoV-E4. The *in vivo* acute and sub-acute toxicity study of ATRICOV 452 did not show any significant toxicity. Further, *in vivo* Anti-SARS-CoV-2 activity needs to be demonstrated and worth investigating the test samples in clinical trials.

## Acknowledgement

**Competing interests:** Latha Damle is the founder of Atrimed Biotech LLP and holds equity in Atrimed Pharmaceuticals. Shiban Ganju and Hrishikesh Damle hold equity shares in Atrimed Pharmaceuticals.

**Grant information:** The study was sponsored by Atrimed Pharmaceuticals Pvt. Ltd and no external funding was received for the study.

